# α-synucleinopathy associated calcium overload and autophagy failure is regulated by gain-of-function of Tousled-like kinase

**DOI:** 10.1101/2022.07.17.500360

**Authors:** Fangyan Gong, Hao Chen, Ying Xiong, Rong Cai, Jing Zhang, Ningli Wang, Lei Liu

## Abstract

As a pathological hallmark in Parkinson’s disease (PD), α-synucleinopathy causes multiple cellular damages, including calcium overload, mitochondrial and autophagic dysfunction, and eventually dopamine neuron death. However, the hierarchy of these detrimental events is unclear. In *Drosophila*, we confirmed that overexpression of α-synuclein could induce all these cytotoxic events. To determine the specific cytotoxic events induced by calcium overload, we established a calcium overload model in *Drosophila* and performed genetic screens. We found that calcium overload caused mitochondrial damage and autophagy failure and cell death, and these cytotoxic processes could be strongly rescued by loss of Tousled-like kinase (TLK). Interestingly, loss of TLK also rescued defects induced by α-synuclein overexpression in *Drosophila*. This suggests that calcium overload acts as the crucial event upstream of mitochondrial and autophagy dysfunction. For TLK regulation of autophagy, our data indicated that a transcriptional factor REPTOR, which regulated the expression of several lysosomal genes, functioned downstream of TLK. In mammalian cells and mice, TLK2 (the homolog of *Drosophila* TLK) was phosphorylated under calcium overload. Upon phosphorylation, TLK2 increased its kinase activity. In addition, TLK2 could phosphorylate CREBRF (the human homolog of REPTOR) to cause its loss of transcription on the lysosomal genes. Moreover, TLK2 knockout mice rescued multi-aspect cytotoxicity induced by calcium overload and α-synuclein overexpression. Our research demonstrates that TLK2 acts as a key regulator to mediate cell death and dysfunctions of mitochondria and autophagy downstream of calcium overload.

## Introduction

Parkinson’s disease (PD) is a common neurodegenerative disorder affecting 2-3% of people over 65 years of age. The prominent pathological hallmark in the brain of PD patients comprise in α-synuclein in dopamine (DA) neurons ^1^. α-synuclein causes multi-aspect of pathological damages, including aberrant protein aggregates, disruption of autophagy-lysosome catabolism, mitochondrial damage and disruption of calcium homeostasis ^2^. Recent studies provide compelling evidence suggesting that loss of calcium homeostasis plays a prominent pathological role in PD ^3^. For instance, the pacemaking activity in DA neurons leads to sustained Ca^2+^ influx through CaV1.3, a L-type voltage-dependent Ca^2+^ channel, which renders neuron vulnerable to stress ^4^. Blocking CaV1.3 by isradipine, a dihydropyridine blocker approved for hypertension treatment, had protective effect in animal models of PD, although it failed a clinical trial ^5^. In fact, intracellular Ca^2+^ elevation may also initiate by a variety of cation channels on cytoplasmic, ER or mitochondrial membranes. Moreover, both extracellular and intracellular α-synuclein oligomers can form Ca^2+^ permeable pores to elevate intracellular Ca^2+^ concentration and cell death ^6, 7^. In addition, α-synuclein fibers may alter packaging of lipid membranes to disrupt cytosolic calcium homeostasis ^8^. Alternatively, α-synuclein oligomers may provoke inositol 1,4,5-triphosphate receptor (IP3R) in ER, impair mitochondrial respiratory complex I function and lysosome, contributing cytosolic Ca^2+^ overload ^3, 9^. Compelling evidence implicates that calcium overload plays a key detrimental role to induce and propagate the pathogenesis of PD; and cathepsin D and calcineurin are known regulators downstream of calcium overload under α-synuclein expression; and partial inhibition of calcineurin was beneficial for neuron survival ^10, 11^. Weather other the key regulator exist downstream of calcium overload is unknown.

Besides calcium overload, most PD-related genes, such as α-synuclein, PINK1, Parkin, LRRK2, DJ-1, GBA, and ATPA13A2, involve in mitochondrial function and autophagy-lysosome pathways; and mutations in these genes causes diverse alternations related to the progression of PD ^12, 13^. Drugs targeting mitochondrial function or autophagy-lysosome pathway have shown a wide-spectrum of protective effects in animal models of PD, however they all failed to slow the progression of PD in patients ^12, 14^.

Among the detrimental mechanism of calcium overload, mitochondrial dysfunction and autophagy-lysosome deficiency, which one is the initial inducer? Whether there exists a crucial regulator to initiate these damages? To answer these questions, we generated a *Drosophila* model of calcium overload and performed genetic screens to discover genetic modifiers. Then, we tested the modifiers in an α-synuclein overexpression induced PD model in *Drosophila*. The results showed that calcium overload caused cell death associated with both mitochondrial and autophagy-lysosome dysfunctions. Strikingly, loss-of-function of Tousled-like kinase (TLK) abolished both calcium-overload-induced and α-synuclein-induced cytotoxicity, suggesting TLK is a key regulator of multi-aspect pathological events in PD. Using *in vitro* biochemical analysis and mouse genetics, we further determined that calcium overload could induce hyper-phosphorylation of TLK2, the mammalian homolog of *Drosophila* TLK. We propose that phosphorylated TLK2 is a key event to induce cytotoxicity after calcium overload.

## Results

### Calcium overload in *Drosophila* induces diverse cytotoxicity and cell death

The cytotoxicity induced by α-synuclein shares many features with calcium overload induced damage, including mitochondrial dysfunction, autophagy-lysosome defect, ER stress, and cell death ^3, 15^. In fact, α-synuclein can induce calcium overload by diverse mechanisms ^6, 7^. To test whether calcium overload plays a crucial detrimental role, we generated a transgene to express a mutant (G430C) of human brain sodium channel 1 (BNC1, also known as ASIC2a), which forms a constitutively opened none-selective cation channel in heterologous expression systems ^16^. Therefore, expression of BNC1_G430C_ is more likely to induce calcium toxicity over sodium toxicity. We generated a BNC1_G430C_ transgenic fly simplified as *UAS-C16* ^17^. For genetic screens, we tested different promoters and observed that *Mhc-gal4* (a muscle promoter) driven *C16* (*Mhc-gal4>C16*, or *MC16*) resulted in a stereotypical wing posture defects in adult flies, a phenotype easy to quantify (Fig. 1A). Crossed with *UAS-GFP*, the muscle damage was observable (Fig. 1B). Using cell permeable dye Fluo4-AM and Fura2-AM, which are calcium indicators, we confirmed that the cytoplasmic Ca^2+^ level of *MC16* flies was increased compared to the control (Fig. 1C and S1A). Indicating *MC16* was indeed a model of calcium overload. Meanwhile, the lifespan of *MC16* flies was shortened (Fig. S1B).

**Figure 1.**
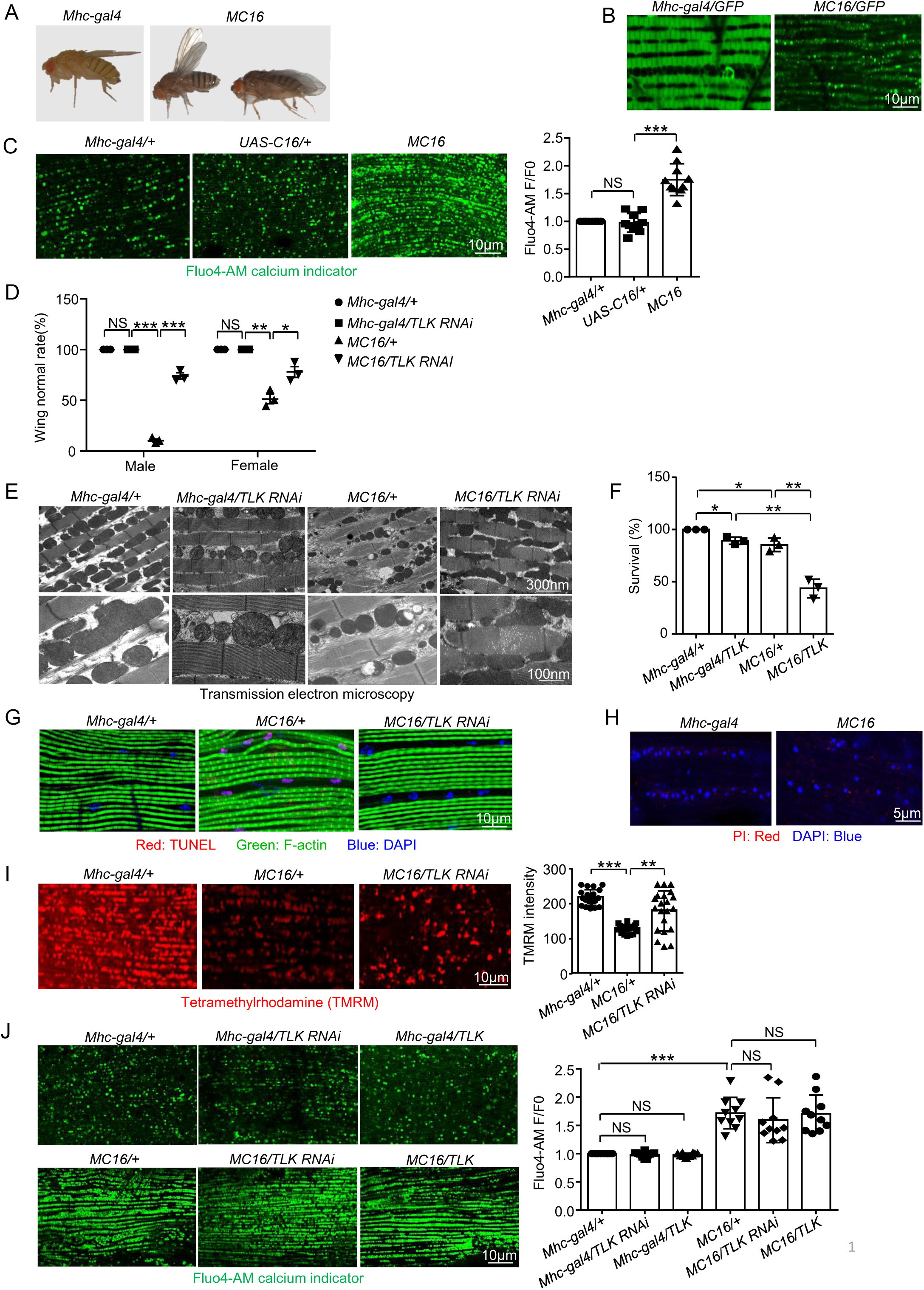
*MC16* is a calcium overload model in *Drosophila* and *TLK RNAi* reduces the diverse cytotoxicity of *MC16* fly. (**A**) The wing phenotype of *MC16* fly. (**B**) Live imaging of the indirect flight muscle morphology of adult flies. The muscles were labeled with *UAS-GFP* driven by *Mhc-gal4.* Scale bar, 10 μm. N=10. (**C**) Calcium overload level was measured by Fluo4-AM (5 μM, green) calcium indicator, the relative intensity was analyzed by F/F0, F represented the fluorescence intensity. Scale bar, 10 μm. N=10. Error bars indicate SD. Statistical significance was performed with an unpaired t-test between two groups. For comparison of more than two groups, significance was determined using a one-way ANOVA with Tukey’s post hoc test. Kaplan-Meier tests were used under the log-rank algorithm for survival analysis. * P < 0.05, ** P < 0.01, *** P < 0.001 and ****p < 0.0001. (**D**) The wing normal rate of the *Mhc-gal4/+, Mhc-gal4/TLK RNAi, MC16/+* and *MC16/TLK RNAi* flies. Both male and female flies were compared; 100 flies were tested for each trial (N=3). (**E**) Transmission electron microscopy (TEM) images of the indirect flight muscle of adult flies. Scale bar, 300 nm and 100 nm. N=5. (**F**) The survival rate of 3-7 days of *Mhc-gal4/+, Mhc-gal4/TLK, MC16/+* and *MC16/TLK* flies were analyzed. 100 flies were tested for each trial (N=3). (**G**) TUNEL (red) staining of the indirect flight muscle, muscles were stained with F-actin (green). Nuclei were stained with DAPI (blue). Scale bar, 10 μm. 6 flies were tested for each trial (N=3). (**H**) Propidium Iodide (PI) staining of the *Mhc-gal4* and *MC16* flies, nuclei were stained with DAPI (blue). Scale bar, 5 μm. 6 flies were tested for each trial (N=3). (**I**) Tetramethylrhodamine (TMRM, red) staining of the indirect flight muscle. The fluorescence intensity was analyzed. Scale bar, 10 μm. N=20. (**J**) The same as **C** with genotype listed on the micrographs. The full gels are shown in the supplementary Figure S1 source data.

### TLK is a key regulator of calcium overload in *Drosophila*

To screen for genetic modifiers against the calcium overload-mediated cell damage, the wing posture defects were quantified. For unknown reason, we observed that the *MC16* male flies displayed a higher percentage of wing defect than the female flies (Fig. 1D). We screened about 437 UAS-RNAi and candidate-based lines (Table S1) and identified genetic modifiers of the following genes: Raptor, CalpB, MED7, MED19, MED20, MED27, TLK, Crtc, Atg8a and Reptor. The RNAi of Crtc, Atg8a and Reptor genes were enhancers; and the RNAi of Raptor, CalpB, MED7, MED19, MED20, MED27 and TLK were suppressors (Table S2). Among them, we found Tousled-like kinase (TLK) RNAi showed the most striking rescue effect (Fig. 1D). Because many RNAi can have off-targets, therefore we used two TLK RNAi lines each targeting different TLK mRNA sequence. By qRT-PCR, their TLK RNAi effects on TLK transcription were confirmed; and they did not affect the expression of *C16* (Fig. S1C and S1D). In addition, TLK RNAi could rescue the muscle morphology and mitochondrial defects of the *MC16* fly (Fig. 1E). Interestingly, overexpression TLK could enhance *MC16* lethality (Fig. 1F), suggesting TLK was necessary and sufficient to regulate calcium overload-induced cell death. We also determined cell death in the *MC16* fly by TdT-mediated dUTP nick end labelling (TUNEL) and propidium iodide (PI), which detects apoptosis and necrosis respectively. Compared to the negative staining in the control flies, TUNEL signals were presented in the muscle cells of *MC16* flies, and TLK RNAi strongly prohibited the cell death (Fig. 1G and S1E). PI staining was negative in the *MC16* flies, indicating the plasma membrane was intact (Fig. 1H). TMRM (Tetramethylrhodamin) is a fluorescent dye that can be sequestered by active mitochondria; its positive staining indicates healthy mitochondria. Similarly, MitoBlue is a membrane potential-independent dye and stains functional mitochondria. Compared to the control flies, fluorescence of TMRM and MitoBlue was greatly reduced in the *MC16* flies, and this defect was restored by TLK RNAi (Fig. 1I and S1F). TLK RNAi and TLK overexpression did not affect calcium overload (Fig. 1J), suggesting TLK functions downstream of calcium overload.

### Cellular damage induced by α-synuclein overexpression is rescued by TLK RNAi in *Drosophila*

To test whether overexpression of α-synuclein induces calcium overload, we obtained a *Drosophila* transgenic line to express α-synuclein. The *UAS-SNCA* (the α-synuclein gene) was driven by a gene switch inducible promotor *daughterless-gal4* (*DaGS*) ^18^. The progeny flies are simplified as *DaGS>SNCA*. Adult flies were fed with RU486 (Mifepristone) to induce the Gal4 expression ubiquitously. Because RU486 did not affect fly health ^19^, *DaGS>SNCA* flies fed with RU486 was compared with the control (*DaGS>SNCA* flies without feeding RU486). In 20 days-old of *DaGS>SNCA* flies, α-synuclein could not induce the calcium overload (Fig. S1G), indicating α-synuclein might not reach the threshold level. In these flies, TLK RNAi could not reduce α-synuclein level and restore the TH (Tyrosine Hydroxylase) level, a marker of DA neuron (Fig. S1H). In 30 days-old *DaGS>SNCA* flies, intracellular Ca^2+^ level was increased in brain and muscle cells (Fig. 2A), indicating α-synuclein indeed promoted calcium overload. The expression of α-synuclein was detectable, associated with reduced TH level (Fig. 2B), suggesting α-synuclein overexpression causes DA neuron death. Importantly, TLK RNAi could reduce the α-synuclein protein level and restore the TH protein level in the *DaGS>SNCA* flies (Fig. 2B). TH immunostaining also showed rescue effects (Fig. 2C and S1J). These results indicate that TLK RNAi maybe effective after TLK activation, which is induced via calcium overload by α-synuclein expression. In addition, we found that TLK overexpression further shortened the lifespan of flies expressing α-synuclein (Fig. 2D). Furthermore, defective mitochondrial function in the *DaGS>SNCA* flies was also restored by TLK RNAi (Fig. 2E); and TLK RNAi did not affect calcium overload induced by α-synuclein expression (Fig. 2F). These results suggest that α-synuclein overexpression causes damage in DA neurons mainly through calcium overload.

**Figure 2.**
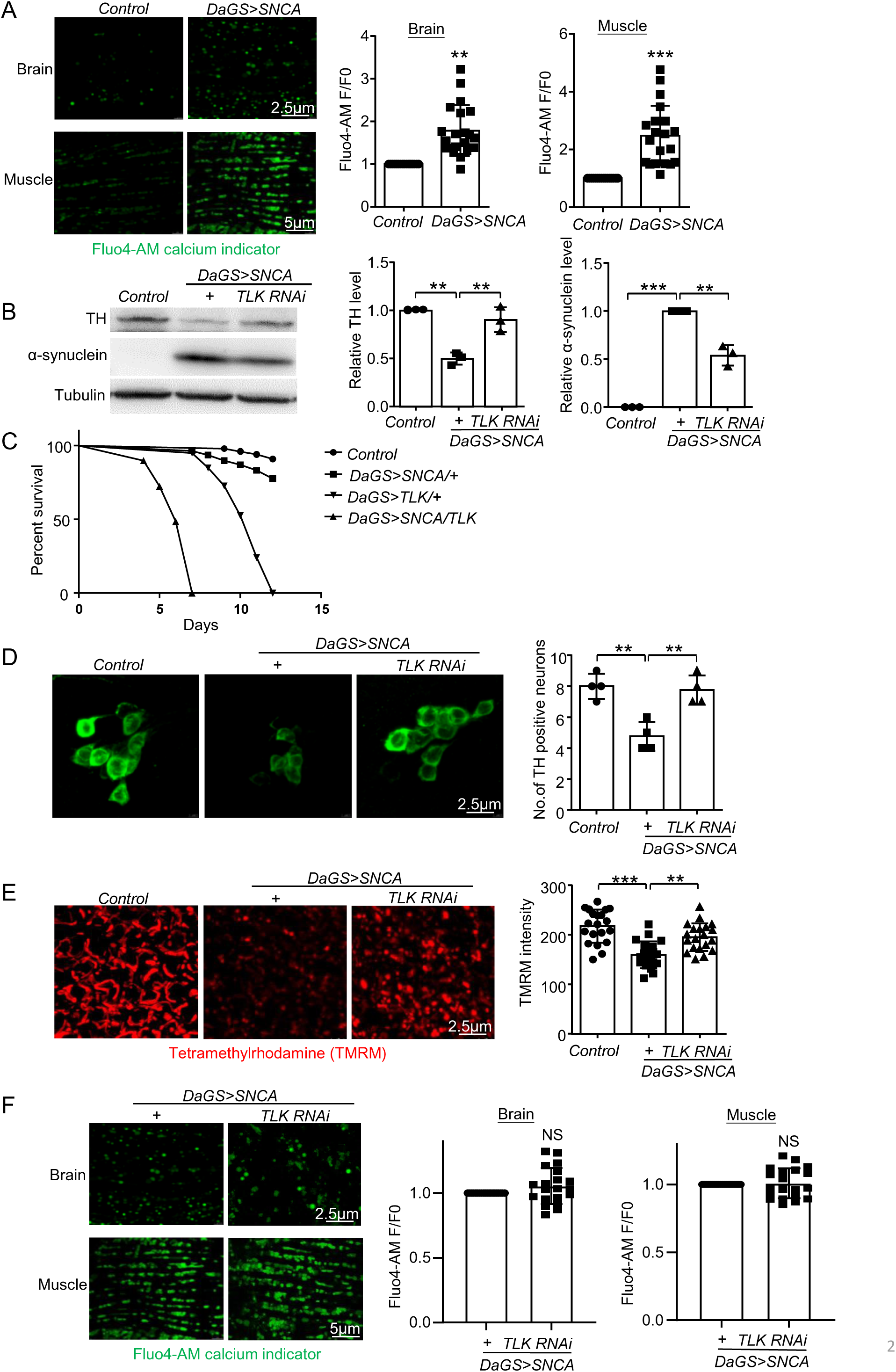
*TLK RNAi* rescues the α-synuclein overexpression induced cellular damage in *Drosophila*. (**A**) Calcium overload level was measured by Fluo4-AM (5 μM, green) calcium indicator of the brain and the muscle in 30 days-old, the relative intensity was analyzed by F/F0, F represented the fluorescence intensity. Brain Scale bar, 2.5 μm, Muscle Scale bar, 5 μm. N=20. (**B**) Western blot of the tissue extracts from the whole body in 30 days-old. Quantification of TH and α-synuclein level. 20 flies were tested for each trial (N=3). (**C**) Immunostaining images of the TH neurons (green) in *DaGS>SNCA* fly brain PPL1 cluster. Scale bar, 2.5 μm. 5 flies were tested for each trial (N=4). Quantification of the number of the TH positive neurons. (**D**) The lifespan changes of *Control* (No RU486), *DaGS>SNCA/+*, *DaGS>TLK/+* and *DaGS>SNCA/TLK* flies. 80-100 flies were tested for each trial (N=3). (E) TMRM (1 uM, red) staining of the fly brain, the fluorescence intensity was analyzed. Scale bar, 2.5 μm. N=20. (**F**) The same as **A** with genotype listed on the micrographs. The full gels are shown in the supplementary Figure 2 source data.

### Calcium overload disrupted lysosome-autophagy pathway via TLK function

To study the mechanism of TLK, we performed whole genome RNA sequencing to compare transcriptional alterations of the control (*Mhc-gal4*) and calcium overload (*MC16*) flies. The volcano plot displayed gene expression difference between the two groups of flies (Fig. S2A); 561 genes upregulated and 924 genes downregulated in the *MC16* flies (Fig. S2A). Compared the *MC16* flies, *TLK RNAi* under *MC16* background resulted in 616 genes upregulated and 531 genes downregulated (Fig. S2B). Compared the 924 downregulated genes in *MC16* and 616 upregulated gene in *MC16/TLK RNAi*, 73 genes were overlapped; and their expression intensity was plotted (Fig. S2C). To further characterize the 73 genes by KEGG (Kyoto Encyclopedia of Genes and Genomes), the pathways of lysosome and protein processing in ER were enriched (Fig. S2D). Using fluorescent markers of *UAS-ER-RFP* (label ER) and mitochondrial marker *UAS-mitoRFP* (label mitochondria), we found that mitochondrial morphology was disrupted and ER morphology showed less change (Fig. S2E and S2F). Therefore, we further focused on the lysosome pathway. First, we performed qRT-PCR to quantify the transcripts of five lysosomal genes identified from the RNAseq. We found that TLK RNAi could affect several lysosomal functional genes, including lysosomal acidification (CG7997, CtsB and Gba1b) and hydrolyzation (VhaSFD) genes with not change of Lamp1 gene. This result indicates that TLK is more likely to affect the lysosome function rather than its biogenesis (Fig. S2G). Next, we examined lysosome-autophagy status. As marker of autophagy, LC3 protein complex involves in the elongation of phagophore membrane; and GABARAP protein is essential for autophagosome maturation ^20^. In addition, Ref(2)P is the *Drosophila* homologue of mammalian P62, which functions as a substrate of autophagy to deliver ubiquitinated protein into autophagosome ^21^. Accumulation of P62 indicates less active autophagy ^22^. In *MC16* flies, Ref(2)P protein level was increased and GABARAP level was unaltered (Fig. 3A). Because TLK mainly affected the lysosomal function, and P62 is a marker of lysosomal degradation. Therefore, P62 protein level changes without GABARAP level. Importantly, TLK RNAi could rescue the autophagic failure (Fig. 3A). Moreover, *MC16* flies decreased the LysoTracker staining and rescued by TLK RNAi (Fig. 3B). This result further supports that lysosome activity is downregulated and this reduction depends on TLK function. Similarly, we observed that the Ref(2)P protein level was increased in the RU486 fed *DaGS>SNCA*, associated with decreased LysoTracker staining (Fig. 3C and 3D); and these deficits could also be rescued by TLK RNAi (Fig. 3C and 3D). Collectively, our results demonstrated that TLK RNAi could rescue the lysosome-autophagy dysfunction downstream of calcium overload or α-synuclein overexpression in *Drosophila*.

**Figure 3.**
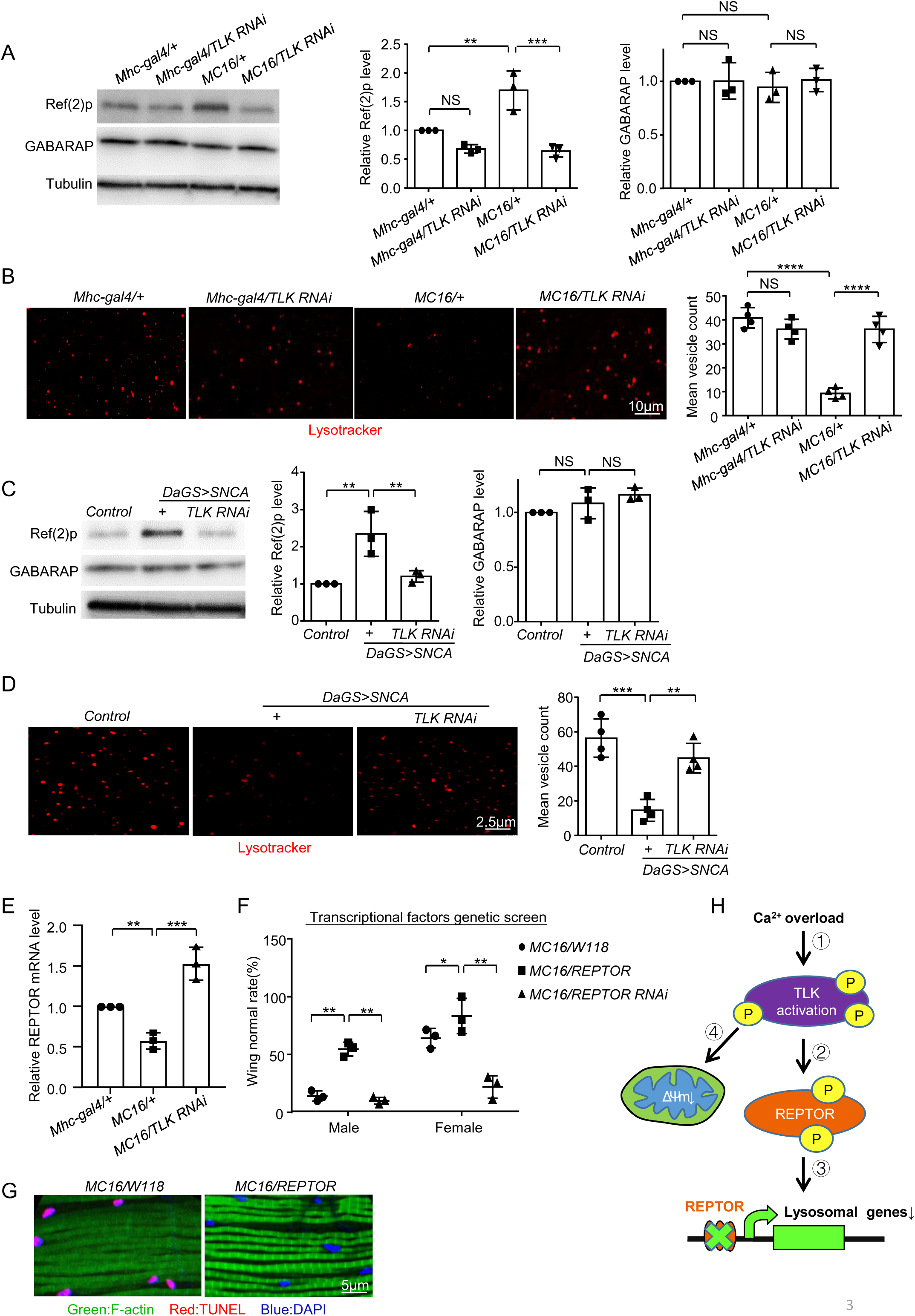
Calcium overload disrupted lysosome-autophagy pathway via TLK function. (**A**) Western blot of the tissue extracts from the indirect flight muscle. 20 flies were tested for each trial (N=3). Quantification of Ref(2)p and GABARAP level. (**B**) Lysotracker (red) staining of the indirect flight muscle. Scale bar, 10 μm. 5 flies were tested for each trial (N=4). Quantification of the number of lysotracker positive vesicles. (**C**) Western blot of the tissue extracts from the whole body. 20 flies were tested for each trial (N=3). Quantification of Ref(2)p and GABARAP level. (**D**) Lysotracker (red) staining of the fly brain. Scale bar, 2.5 μm. 5 flies were tested for each trial (N=4). Quantification of the number of lysotracker positive vesicles. (**E**) The wing normal rate of 3-7 days flies were analyzed after transcriptional factors genetic screen. 50−60 flies were tested for each trial (N=3). (**F**) TUNEL (red) staining of the indirect flight muscle, muscles were stained with F-actin (green). Scale bar, 5 μm. 5 flies were tested for each trial (N=3). (**G**) A schematic model of calcium overload cytotoxicity. The full gels are shown in the supplementary Figure 3 and Figure S3 source data.

How TLK may regulate lysosome-autophagy function? It is known that TLK is a serine/threonine kinase to function in transcription and DNA repair ^23^. Because transcription of several lysosomal genes was reduced in *MC16* flies and rescued by *TLK RNAi,* we proposed that TLK might affect a transcription factor to suppress the expression of lysosomal genes. Through genetic screens, we identified several transcriptional factors. We suspect that some of the lines might affect C16 expression, especially the MED family genes. Among them, REPTOR RNAi did not affect C16 expression (Fig. S1D). REPTOR is known as a transcription factor responding to the TORC1 signaling to regulate expression of lysosome-autophagy genes ^24^. Functionally, REPTOR RNAi enhanced and REPTOR overexpression rescued the wing defects and cell death of *MC16* flies (Fig. 3E and 3F). Collectively, the fly research suggests the following hypothesis: 1) Calcium overload activates TLK; 2) Activated TLK phosphorylates REPTOR; 3) phosphorylated REPTOR decreases its transcription control on several lysosomal genes. 4) Activated TLK causes mitochondrial dysfunction through an unknown mechanism (Fig. 3G). In the following, we test this hypothesis in mammalian cells and mice.

### Knockout TLK2 activates lysosome-autophagy function

Vertebrate genomes encode two *Drosophila* TLK homologs, TLK1 and TLK2 ^25^. Using CRISPR/Cas9 system, we knocked out human TLK1 and TLK2 genes in Hela cells, respectively. To determine their effect on autophagy, we quantified the transcriptional change of five lysosomal genes (GLA, CTSB, ATP6V1H, GBA and LAMP1) by qRT-PCR. The results showed that TLK2 KO, but not TLK1 KO, increased transcription of three genes (GLA, CTSB and ATP6V1H) (Fig. S3A and S3B), indicating TLK2, but not TLK1 regulates lysosomal function. After induction of autophagy in Earle’s starvation buffer (EBSS) ^26^, these transcripts were further increased (Fig. S3A). In the TLK2 KO cells, the ratio of LC3-II/LC3-1was increased associated with decreased P62 level, but not TLK1 KO (Fig. 4A and S3C), suggesting that TLK2 KO activated autophagy. As a dynamic process, autophagic flux can be assessed by drug treatment to stop the flux at certain stage. Bafilomycin A1 (Baf A1) inhibits the fusion of autophagosomes to lysosomes; rapamycin (Rapa) activates autophagy ^26^. In wild type Hela cells, Rapa treatment increased the LC3-II/LC3-1 ratio and decreased the P62 level; and Baf A1 treatment increased the LC3-II/LC3-1 ratio and P62 level (Fig. 4A). In the TLK2 KO cells, Rapa treatment further activated autophagy by decreasing P62 level, and the activated autophagy was abolished by Baf A1 treatment (Fig. 4A). Furthermore, the transmission electron microscopy (TEM) imaging showed that more autophagosome and autolysosome were present in the TLK2 KO cells than the wild type cells (Fig. 4B). Another commonly used assay to measure autophagic flux is LC3-mRFP-GFP adenovirus infection ^26^. The low pH inside the lysosome quenches the GFP fluorescent signal of LC3-mRFP-GFP; in contrast, RFP exhibits more stable fluorescence in acidic compartments. If autophagic flux is increased, both yellow and red puncta will increase. The yellow stains autophagosome and the red labels autolysosome. Indeed, both yellow and red puncta were increased in the TLK2 KO cells (Fig. 4C), suggesting increased autophagic flux. As additional support, P62 protein level was lower in P62 overexpression in TLK2 KO cells (Fig. S3D). These results are consistent with a previous study of a genome-wide siRNA screen, in which siRNA TLK2 enhanced the amino acid starvation-induced autophagy^27^. Together, these results demonstrated that TLK2 KO activated lysosome-autophagy function.

**Figure 4.**
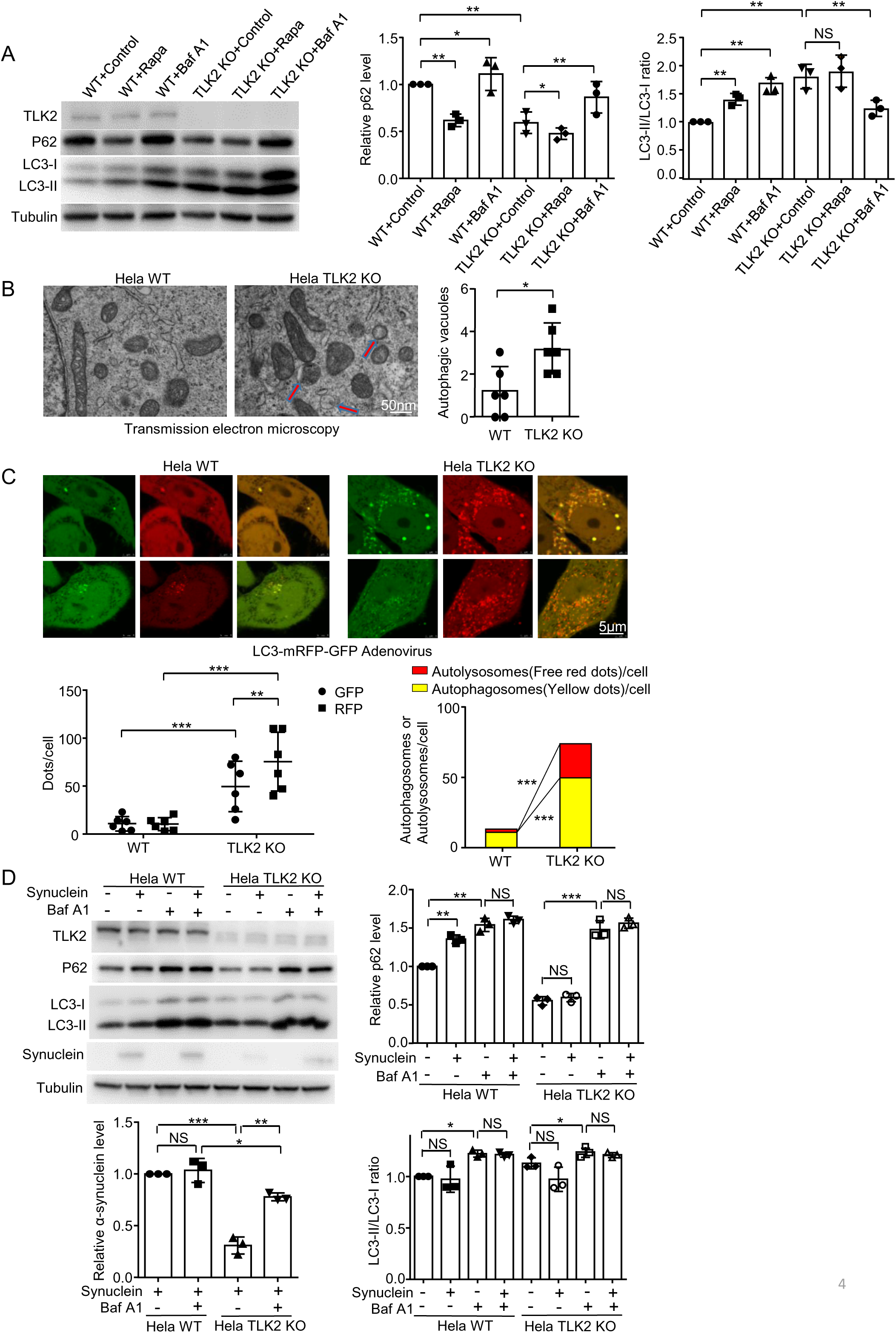
Knockout TLK2 activates lysosome-autophagy function. (**A**) Western blot of the extracts from Hela WT and TLK2 KO cells with Rapamycin (Rapa; 40 nM for 6 hours) and Bafilomycin A1 (BafA1, 100 nM for 6 hours) treatment. N=3. Quantification of P62 and LC3-II/LC3-I ratio level. (**B**) TEM images of the Hela WT and TLK2 KO cells. Scale bar, 50 nm. N=6. Quantification of the number of autophagic vacuoles (red arrows). (**C**) Live imaging of the Hela WT and TLK2 KO cells after LC3-mRFP-GFP adenovirus transfection 48h. Scale bar, 5 μm. N=6. Quantification of the GFP and RFP dots, as well as autolysosomes (free red dots) and autophagosomes (yellow dots) per cell. (**D**) Western blot of the extracts from Hela WT and TLK2 KO cells with Bafilomycin A1 (BafA1, 100 nM for 6 hours) treatment. Using lentiviral transfection of α-synuclein, we established stable cell lines expressing α-synuclein in wild type Hela cells and TLK2 KO cells. N=3. Quantification of α-synuclein, P62, LC3-II/LC3-I ratio and Lamp2a level. The full gels are shown in the supplementary Figure 4 source data.

To study a cellular model of PD, we established stable cell lines expressing α-synuclein in wild type Hela cells and TLK2 KO cells by lentiviral transfection of α-synuclein. Then we treated the cell lines with Bafilomycin A1. The results showed that α-synuclein level decreased significantly under the TLK2 KO background compared with the wild type. Baf A1 treatment restored the α-synuclein level (Fig. 4D). We also detected the LC3 level and P62 level. We found that α-synuclein overexpression could increase the P62 level in the Hela WT cells but not in the TLK2 KO cells (Fig. 4D), consistent with drosophila data. In neuroblastoma SKN-SH cells, α-synuclein overexpression showed a lower LC3-II level and higher P62 level compared to normal cells (Fig. S3E), suggesting α-synuclein overexpression reduced autophagy function. This effect is consistent with reports by others ^28^. Interestingly, the downregulated autophagy could be rescued by a known TLK2 kinase inhibitor, promazine hydrochloride (PMZ) (Fig. S3E). In addition, LysoTracker signal was declined in α-synuclein overexpressed SKN-SH cells; and PMZ rescued the lysosome dysfunction (Fig. S3F). These results indicate that TLK2 may involve in regulation of α-synuclein overexpression induced autophagy failure.

### Calcium overload increased TLK2 phosphorylation; and CREBRF was a substrate of TLK2

TLK2 contains multiple phosphorylation sites and autophosphorylation can activate this enzyme ^29^. To test whether TLK2 was phosphorylated during calcium overload, we constructed GST-TLK2 and pulled down TLK2 protein by GST. We purified TLK2 and incubated with cell lysate from ionomycin treated cells (induced calcium overload) or cell lysate from normal cells. Strikingly, we observed a shift of GST-TLK2 after incubation with ionomycin-treated cell lysate on the Coomassie Blue gel (Fig. 5A), indicating TLK2 protein is modified. In addition, incubated with ionomycin-treated cell lysate from TLK2 KO cells showed a similar shift, suggesting the modification of TLK2 is not by itself (Fig. 5A). By phos-tag SDS-PAGE, we found TLK2 phosphorylation was increased (Fig. 5B). By mass spectrometry, all serine/threonine phosphorylation sites of TLK2 were determined. In normal condition (without ionomycin treatment), 12 sites were phosphorylated, including 10 sites shared in control and ionomycin treated (S102, S114, S116, S133, T160, S222, S329, S375, S392 and S449); and 2 unique sites in control (S225 and S376); while treated with calcium overload lysate resulted in 15 more phosphorylation sites (S25, S72, S109, S110, T207, S209, T212, S217, S277, T299, T300, T366, S616, S751 and S752), associated with loss of 2 phosphorylation sites (S225 and S376) (Fig. 5C).

**Figure 5.**
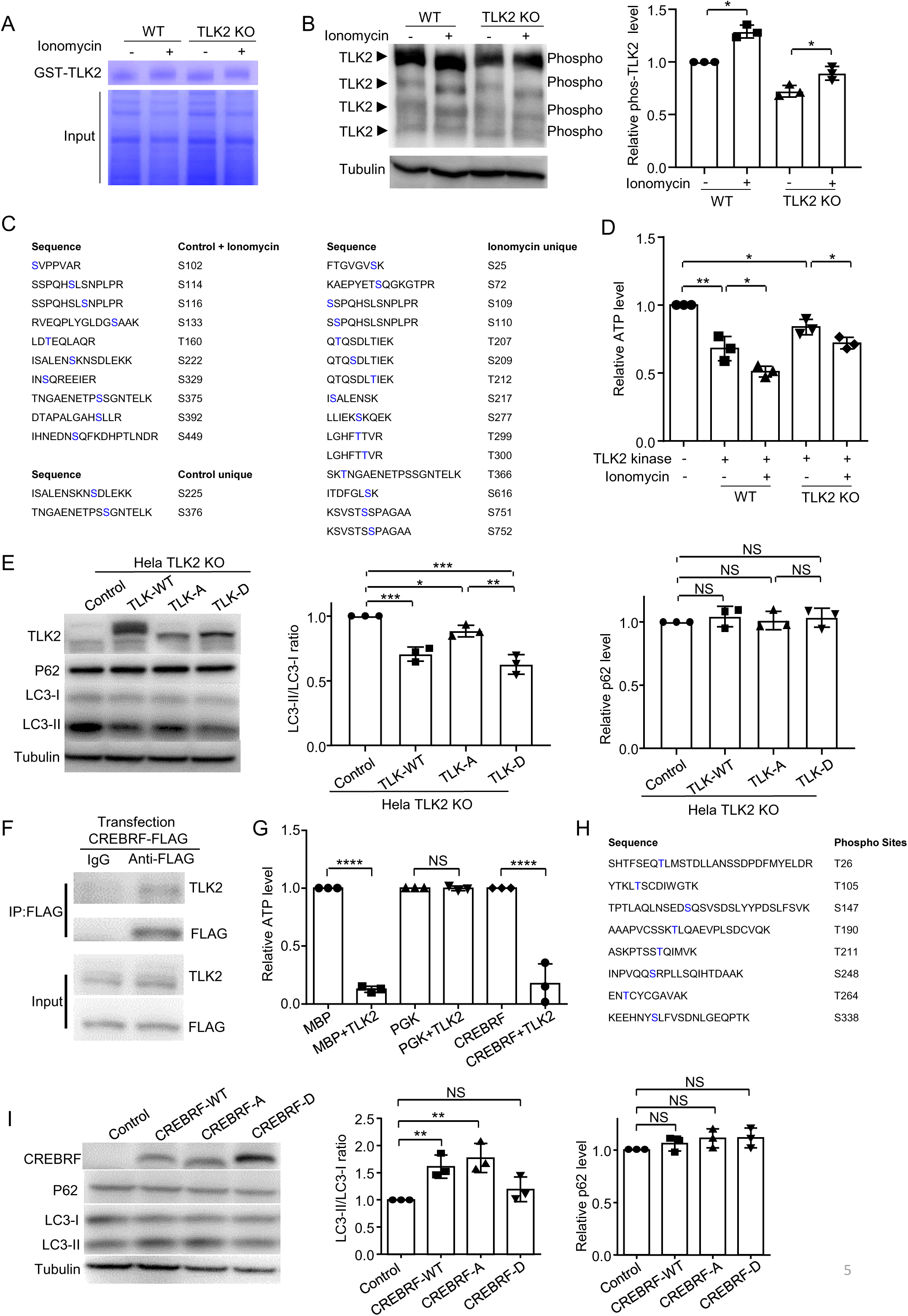
Calcium overload increased TLK2 phosphorylation and CREBRF was a substrate of TLK2. (**A**) Coomassie Blue staining of the GST-TLK2 pull-down from the Hela WT and TLK2 KO cells with or without ionomycin (1 μM) treatment. N=3. (**B**) Phos-tag western blot of the phosphorylated GST-TLK2. Hela WT and TLK2 KO cells with or without ionomycin (1 μM) treatment. GST-TLK2 purification protein were incubated with the cell lysate and pulled down by Glutathione Sepharose 4B beads. Tubulin was used as a loading control. N=3. Quantification of TLK2 phosphorylation level. (**C**) Mass spectrometry results showed the phosphorylation sites after the incubation. (**D**) In vitro kinase assay, recombinant TLK2 protein in *E. coli* were purified. Myelin basic protein (MBP) was used as a substrate. The TLK2 kinase activity with or without ionomycin treatment were assessed by the reduced ATP level. N=3. (**E**) Western blot of the Hela TLK2 KO cell extracts after 48h transfection of TLK2-WT (normal), TLK2-A (dephosphorylation), TLK2-D (phosphorylation) and control. N=3. Quantification of P62 and LC3-II/LC3-I ratio level. (**F**) Co-IP of CREBRF-FLAG and TLK2. CREBRF-FLAG was transfected into HEK293T cells for 48h. Anti-flag antibody was used to pull down protein from the cell lysate. N=3. (**G** and **H**) TLK2 kinase were tested for different substrate. MBP was a positive control, PGK1 was a negative control, CREBRF was reaction with TLK2. N=3. Phosphorylation sites were recognized by mass spectrometry after the reaction. (**I**) Western blot of the cell extracts after 48 hours transfection of CREBRF-WT (normal), CREBRF-A (dephosphorylation), CREBRF-D (phosphorylation) and control. N=3. Quantification of P62 and LC3-II/LC3-I ratio level. The full gels are shown in the supplementary Figure 5 source data.

For the kinase assay *in vitro*, recombinant TLK2 protein in *E. coli* were purified. Myelin basic protein (MBP) has used as a substrate for many kinases ^30^. In this kinase assay, calcium-overload-lysate-treated TLK2 resulted in an increased activity under both WT and TLK2 KO lysates, determined by consumption of ATP (Fig. 5D). However, The TLK2 KO lysates showed less ATP decline compared to the WT lysates (Fig. 5D), indicating TLK2 may phosphorylate itself. This result is consistent with a previous report suggesting that phosphorylation of TLK2 enhances its enzymatic activity ^29^. Moreover, other unknown kinase(s) was activated by calcium overload and involved in phosphorylation of TLK2. By mass spectrometry, the phosphorylation of TLK2 occurred at 15 sites, including S25, S72, S109, S110, T207, S209, T212, S217, S277, T299, T300, T366, S616, S751 and S752. We mutated these 15 sites to either alanine (A) to mimic dephosphorylation or aspartic acid (D) to mimic phosphorylation, and expressed them in both wild type and TLK2 KO Hela cells. The results showed that overexpression of TLK2-WT or TLK2-D decreased the LC3-II/LC3-I ratio; but not overexpression of TLK2-A (Fig. 5E), indicating the phosphorylated form of TLK2 suppresses autophagy. The TLK2-WT migrating differently compared to TLK2-D and TLK2-A on the gel. It is likely that the TLK2-A and TLK2-D mutations may affect phosphorylation of TLK2 at the other sites, as under normal condition TLK2 had 12 phosphorylation sites (Fig. 5C). With more phosphorylation sites, migration of TLK2-WT on the gel may be slower.

The mammalian homolog of *Drosophila* REPTOR is CREB3 regulatory factor (CREBRF) ^24^. We found that TLK2 could interact with CREBRF-flag by co-immunoprecipitation (co-IP) in 293T cells (Fig. 5F). The *in vitro* kinase assay showed that CREBRF was indeed a substrate of TLK2 (Fig. 5G). By mass spectrometry, eight sites of CREBRF were phosphorylated (T26, T105, S147, T190, T211, S248, T264 and S338) (Fig. 5H). To test whether CREBRF phosphorylation might affect its transcriptional function, we mutated these serine and threonine sites to either alanine (A) or aspartic acid (D). After overexpression of the CREBRF-WT, CREBRF-A and CREBRF-D in the MES23.5 dopaminergic cells, the transcript levels of lysosomal genes (GLA, CTSB, ATP6V1H, GBA and LAMP1) were quantified by qRT-PCR. The result showed that CREBRF-WT and CREBRF-A overexpression increased levels of GLA, CTSB and ATP6V1H; CREBRF-D overexpression had no effect (Fig. S3G). Further, CREBRF-WT and CREBRF-A overexpression increased the LC3-II/LC3-I ratio, but not the CREBRF-D overexpression (Fig. 5I), suggesting CREBRF phosphorylation reduces autophagy. Taken together, calcium overload triggers TLK2 phosphorylation, which phosphorylates CREBRF to downregulate its transcriptional activity on several lysosomal genes.

### TLK2 KO rescued calcium overload induced DA neuron loss and α-synuclein-induced lesion in mice

To determine TLK2 effect on calcium-overload induced damage *in vivo*, we constructed a calcium overload model in mice. The glutamate receptor 1 Lurcher mutant (GluR1^Lc^) forms constitutively opened cation channel in a heterologous expression system ^31^. We generated a transgenic mouse line which expresses GluR1^Lc^ conditionally (GluR1^Lc^ ^m/m^, “m” for GluR1^Lc^). Crossed with a doxycycline (dox) inducible promoter line (rtTA), the progeny mice (GluR1^Lc^ ^m/+^; rtTA^+/-^) expresses GluR1^Lc^ ubiquitously when doxycycline is provided. We found that providing dox at normal concentration (2 mg/ml) in drinking water induced mouse lethality within one day. After dox dilution of 200 times (0.01 mg/ml), the GluR1^Lc^ mice could survive more than 3 days. Next, we obtained ubiquitous and conditional TLK2 knockout mice (TLK2^flox/flox^; UBC-Cre^ERT2/+^). Crossing these mice, we acquired the desired genotype (GluR1^Lc^ ^m/+^; rtTA^+/-^; TLK2^flox/-^; UBC-Cre^ERT2/+^). The experimental design was outlined (Fig. 6A). Mice with 8-10 weeks of age were administered with tamoxifen (with corn oil as a control) via intraperitoneal injection for 5 days to obtain heterologous TLK2 KO. One month later, dox (0.01 mg/ml) was provided in drinking water to induce the GluR1^Lc^ expression. We found the mice expressed GluR1^Lc^ started to die at day 3 after dox treatment; while mice with heterozygous TLK2 KO survived longer (Fig. 6B and supplementary video). Moreover, both TH level and TH positive neurons were reduced in the GluR1^Lc^ mice; and TLK2 KO alleviated this lesion (Fig. 6C and 6D). Similar as the case in *Drosophila*, the P62 protein level was increased in the GluR1^Lc^ mice; and TLK2 KO could reduce the P62 level (Fig. 6C).

**Figure 6.**
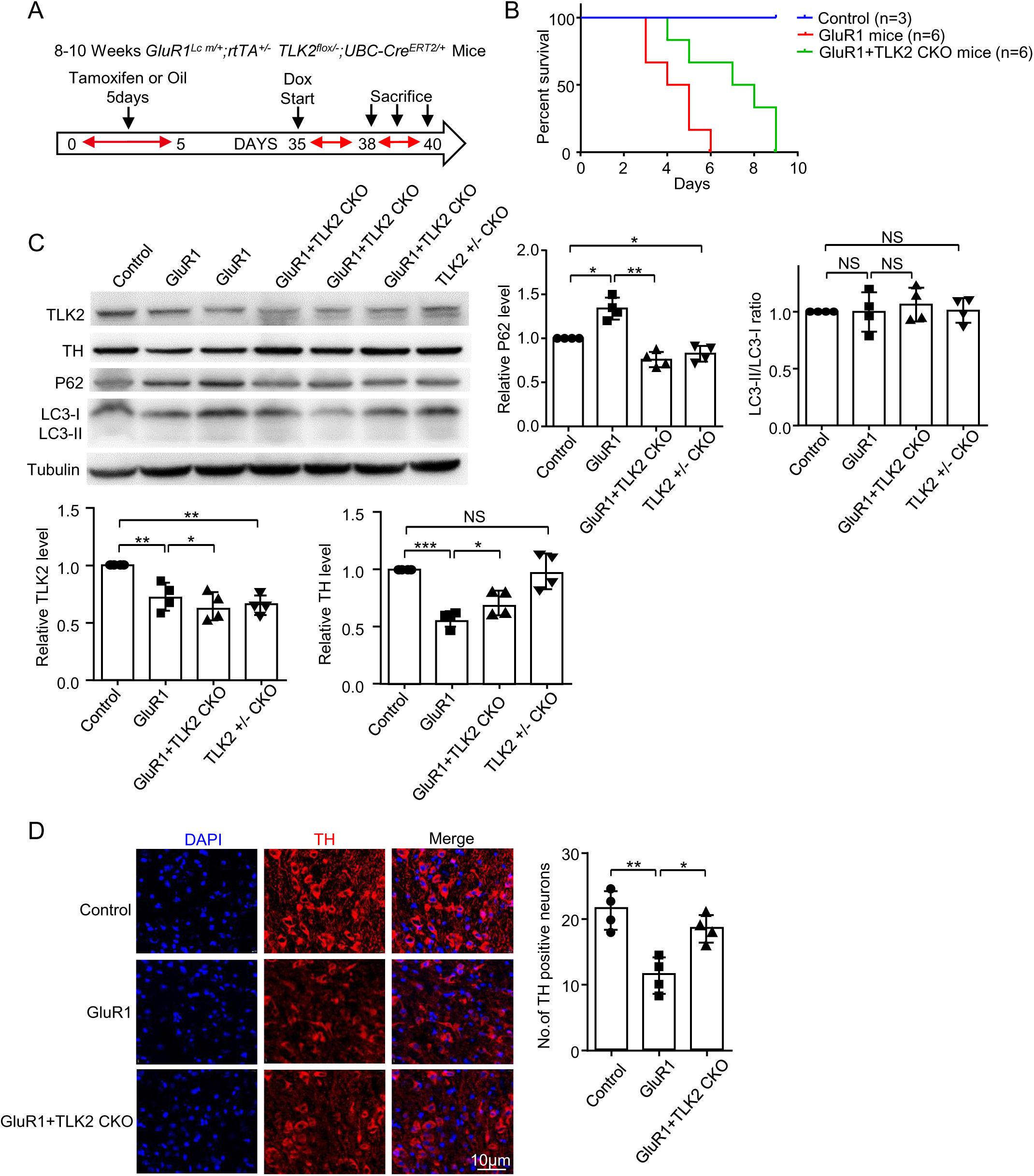
TLK2 KO rescued calcium overload induced DA neuron loss. (**A**) Schematic for experimental design in panels B-D. Mice with 8-10 weeks-old were administered with tamoxifen (corn oil as the control) via intraperitoneal injection for 5 days to obtain heterologous TLK2 KO. One month later, doxycycline (dox) was given in drinking water to induce GluR1^Lc^ expression for 3-5 days. (**B**) Survival rate of the GluR1^Lc^, GluR1^Lc^ / TLK2 CKO and control mice. Control mice (N=3), GluR1^Lc^ and GluR1^Lc^ / TLK2 CKO (N=6). (**C**) Western blot of the tissue extracts from the GluR1^Lc^, GluR1^Lc^ / TLK2 CKO and control mice. N=4. Quantification of TLK2, TH, P62 and LC3-II/LC3-I ratio level. (**D**) Immunostaining images of the TH neurons (red) in the SNc, nuclei were stained with DAPI (blue). Scale bar, 10 μm. N=4. Quantification of the number of the TH positive neurons. The full gels are shown in the supplementary Figure 6 source data.

We also tested the TLK2 KO effect on α-synuclein induced lesion in mice. Injection of adeno-associated virus (AAV) expressed human α-synuclein in mouse midbrain induced a neuropathological lesion in nigral DA neurons, including increased α-synuclein inclusions and progressive axonal degeneration ^32^. The time frame of the experiment was outlined (Fig. 7A). Homozygous TLK2 KO mice (TLK2^flox/flox^; UBC-Cre^ERT2/+^) with 8-10 weeks of age were obtained by tamoxifen administration (with corn oil as a control) via intraperitoneal injection. One week later, these mice were injected with AAV9-CMV-human-α-synuclein or PBS in the right side of substantia nigra pars compacta (SNc). One month after the injection, the mice were tested for behavioral abnormality. In both rotarod and open field tests, the mice expressing α-synuclein performed worse than the control group; and TLK2 CKO mice showed better performance (Fig. S4A and S4B). Compared to the left side of brain (control), the lesion right side of brains showed reduced TH protein level, increased α-synuclein and P62 protein levels; and these defects were alleviated by the TLK2 CKO (Fig. 7B and C). Moreover, the sagittal section of the mouse brain showed the lesion side (the left side of the micrograph) increased α-synuclein accumulation associated with reduced TH positive staining; these defects were alleviated under the TLK2 CKO background (Fig. S4C). Ser129-phosphorylated α-synuclein has been considered as a marker of α-synuclein neuropathology ^33^. Mice overexpressed α-synuclein showed accumulation of Ser129-phosphorylated α-synuclein on the injection side; and TLK2 KO significantly reduced this increase (Fig. 7C). Strikingly, increased TLK2 phosphorylation was detected in the brains of GluR1^Lc^ and α-synuclein overexpression mice (Fig 7E), suggesting TLK2 is activated under these pathological conditions. Collectively, these results indicate that TLK2 plays a key cytotoxic role in α-synucleinopathy.

**Figure 7.**
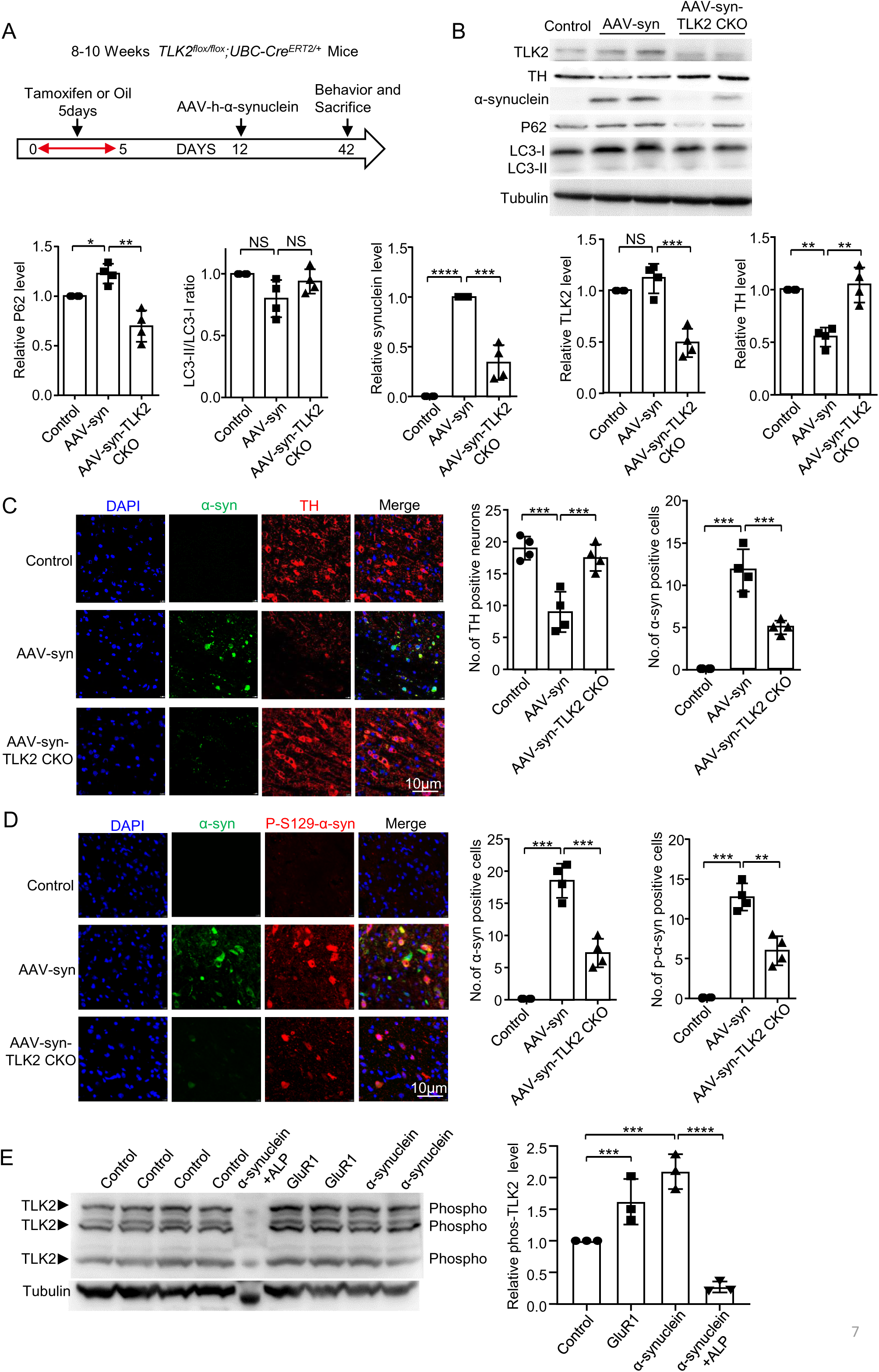
TLK2 KO rescued α-synuclein-induced lesion in mice. (**A**) A schematic experimental design in panels **B**-**F**, homozygous TLK2 KO mice (TLK2^flox/flox^; UBC-Cre^ERT2/+^) at 8-10 weeks-old were obtained by administration tamoxifen (with corn oil as a control) via intraperitoneal injection. One week later, these mice were injected with AAV9-CMV-human-α-synuclein or PBS in the SNc. One month after the injection, the mice were tested for behavioral alternation. (**B**) Western blot of the tissue extracts from the AAV-syn, AAV-syn+TLK2 CKO, and control mice. N=4. Quantification of TLK2, TH, α-synuclein, P62 and LC3-II/LC3-I ratio level. (**C**) Immunostaining images of the TH neurons (red) in the SNc, nuclei were stained with DAPI (blue), α-synuclein were accompanied with GFP fluorescent protein (green). Scale bar, 10 µm. N=4. Quantification of the number of the TH and α-synuclein positive cells. (**D**) Immunostaining images of the P-S129-α-synuclein (red) in the SNc, nuclei were stained with DAPI (blue), α-synuclein were accompanied with GFP fluorescent protein (green). Scale bar, 10 μm. N=4. Quantification of the number of the P-S129-α-synuclein and α-synuclein positive cells. (**E**) Phos-tag western blot of the tissue extracts from the GluR1^Lc^, α-synuclein overexpression and control mice. ALP was used for dephosphorylation of protein serine, threonine and tyrosine residues. Tubulin was used as a loading control. N=3. Quantification of TLK2 phosphorylation level. The full gels are shown in the supplementary Figure 7 source data.

## Discussion

### Disruption of calcium homeostasis is a key cytotoxic event of α-synucleinopathy

Mounting evidence demonstrate that PD-related mutations or environmental risk factors such as aging and exposure of neurotoxins associated with defective calcium homeostasis through a wide variety of mechanisms ^3^. Because calcium signaling plays prominent roles involving multi-aspects of physiology in neurons, loss of calcium homeostasis is clearly pathogenic, especially its effect on mitochondrial dysfunction and disruption of autophagy-lysosome pathway ^34, 35^. Interestingly, selective disruption of mitochondrial function, such as knockout of mitochondrial complex 1 subunit (Ndufs4 or Ndufs2), does not induce significant loss of DA neurons; instead it causes reduction of dopamine content and progressive loss of axons and dendrites of DA neurons ^36, 37^. This indicates that cell death induced by calcium overload is unlikely mediated through disruption of the mitochondrial function.

As a hallmark of PD, accumulation of α-synuclein induces cytotoxicity by diverse mechanisms, including defective calcium homeostasis, ER, Golgi, nuclear, mitochondrial and autophagy-lysosomal functions ^15^. The shared spectrum of cytotoxicity induced by calcium overload and α-synuclein accumulation promoted us to search for their common regulators. Here, we studied a calcium overload model *MC16* in *Drosophila*. After induction of the leaky cation channel, fly muscle cells showed a slow and progressive damage in the first 3-7 days after eclosion. This suggests that *MC16* induces a slow onset of calcium overload, significantly milder than the neuron necrosis model we reported previously ^38^. Consistent with a mild calcium overload stress, the cell death in the muscle cells of *MC16* was not necrosis, because of negative PI staining. In addition, our data demonstrated that α-synuclein overexpression (*DaGS>SNCA*) in *Drosophila* could induce calcium overload, autophagy and mitochondrial dysfunction, and DA neuron death, and these deficits could be rescued by TLK RNAi. These results suggest that calcium overload is the crucial cytotoxic event of α-synucleinopathy.

Although calcium overload is not a cytotoxic event specific to DA neurons, it is likely that DA neurons are prone to calcium overload more than other cell types. Such as both pacemaking activity and expression of specific L-type calcium channels may promote DA neurons to overload calcium ^3^. In addition, α-synuclein presented intracellular or extracellular with monomeric or aggregated form all disrupt calcium homeostasis by diverse mechanisms ^6, 7^. Results from our *Drosophila* models of calcium overload and α-synucleinopathy indicate that calcium overload is at the upstream to cause mitochondrial and autophagic dysfunction. Therefore, these combined stresses may promote eventual cell death. Meanwhile, we cannot rule out that α-synucleinopathy may cause cellular damage on its own, because expression of α-synuclein could reduce the TH level (Fig. 1SH) without calcium overload (Fig. S1G).

### Calcium overload induces TLK2 hyper-phosphorylation which functions to suppress autophagy-lysosome pathway and mitochondrial damage

Mammalian genomes contain two *Drosophila* TLK homolog, TLK1 and TLK2 ^23^. We observed that TLK2 KO (but not TLK1 KO) enhanced autophagy in Hela cells, suggesting TLK2 plays a distinct role from TLK1 to regulate autophagy. For the mechanism, both *Drosophila* TLK and human TLK2 regulated several lysosomal genes at the transcriptional level. In both *MC16* and *DaGS>SNCA* flies, the autophagosomal marker GABARAP-1 did not change but the lysosomal markers were declined (P62 accumulation and less LysoTracker positive staining), indicating defective autolysosomal function. Here, we cannot rule out a deficit of proteasome induced by α-synuclein that causes less P62 degradation, which requires further investigation ^39^.

Interestingly, DA neurons with α-synuclein aggregates also showed reduced lysosomal marker in PD patients ^40^. For the mechanism, our data suggested that calcium overload might regulate autolysosome through transcription. First, several lysosomal transcripts were declined in *MC16* flies; and this reduction could be rescued by TLK RNAi. Second, the expression of *Drosophila* REPTOR, a known transcription factor to regulate lysosome downstream of TORC1 ^24^, was higher under TLK RNAi background and REPTOR overexpression rescued *MC16* defects. Third, our data suggested that CREBRF functioned as a positive regulator in autophagy because overexpression CREBRF increased mRNA level of several lysosomal genes and the LC3-II/LC3-1 ratio in MES23.5 cells. CREBRF has been known as an unfolded protein response-dependent leucine zipper transcription factor, which recruits the nuclear CREB3 out of nucleus and promotes CREB3 protein degradation ^41^. Several studies suggested that CREBRF might act as an inhibitor of autophagy in glioblastoma cells via the CREB3/ATG5 pathway ^42^; while, others reported that CREBRF might function as a positive regulator in autophagy via the mTOR pathway during early pregnancy ^43^, indicating CREBRF may regulate autophagy in a cell type and content specific manner. Our data showed that expression of a phosphorylated CREBRF mutant abolished its effect on transcription of lysosomal genes, suggesting CREBRF phosphorylation by TLK2 eliminates its transcriptional activity. Together, these results suggest that calcium overload is upstream of autolysosome deficiency in α-synucleinopathy. Others reported that calcineurin may regulate lysosomal activity downstream of calcium overload ^10^.

We detected 15 serine/threonine sites of TLK2 to be phosphorylated when purified TLK2 was treated with cell lysate underwent calcium overload. Our data suggest that TLK2 may phosphorylate itself. In addition, an unknown kinase(s) also involves TLK2 phosphorylation under calcium overload condition, which is interesting to investigate in the future. Our data demonstrated that TLK2 phosphorylation occurred in GluR1^Lc^ mice and α-synuclein overexpression mice. Therefore, phosphorylated TLK2 may serve as a marker for cytotoxicity of calcium overload.

### Strategy to target TLK2

Normal functions of TLK1 and TLK2 involve in chromatin assembly, DNA damage response, and transcription ^44, 45^. TLK2 haploinsufficiency causes distinct neurodevelopmental disorders, and TLKs upregulation drives cancer cell proliferation ^46, 47^. In this study, we found that TLK2 heterozygous KO could reduce calcium overload triggered neurodegeneration; and TLK2 conditional homozygous KO could rescue α-synuclein-induced DA neuron death. We observed that TLK2 KO in adult stage did not generate observable behavioral defects in mice, such as appearance, mobility and weight gain, suggesting a safety profile to target TLK2 in the adult stage. In addition, hyper-phosphorylated TLK2 is likely to occur in calcium overload state. Targeting specifically to this TLK2 species may be desirable. Our result showed that promazine hydrochloride (PMZ), a TLK2 kinase inhibitor, could rescue the α-synuclein induced lysosomal decline in SKN-SH cells. However, PMZ showed pronounced side effects as antipsychotic drug, and to be deprescribed ^48^. Finding most specific TLK2 inhibitors are desirable in the future.

### Limitations of this study

We have not determined the threshold α-synuclein level to induced calcium overload. We also don’t know the threshold of calcium overload to trigger cell death in different cell types; and whether TLK2 phosphorylation is sufficient to induce cell death. Moreover, we don’t know the mechanism of cell death induced by *Drosophila* TLK and mammalian TLK2, with the roles of ER and mitochondria in the cell death process. Understanding these questions are important to elucidate the pathology of calcium over in the contest of related neurodegenerative diseases.

In summary, this research demonstrates that calcium overload plays a prominent role for the cytotoxicity of α-synucleinopathy, especially in promotion of cell death. Activation of TLK induces cytotoxicity via silencing of the autolysosomal pathway and induction of mitochondrial damage. The discovery of TLK-mediated signaling pathway provides us a novel target to treat PD and other neurodegenerative conditions with chronic calcium overload.

## Author contributions

Design (FG, NW and LL), data acquisition (FG, HC, YX, RC, CZ and JZ), writing (FG, NW and LL)

## Acknowledgments

This work is supported by grants provide to L.L. by Beijing Natural Science Foundation Program and Scientific Research Key Program of Beijing Municipal Commission of Education (KZ202010025031) and the National Key R&D Program of China (2020YFC2008004).

## Declaration of interests

We declare no interests of any kinds.

## Data availability statement

All data have provided in the submission; and all materials are available upon request.

## Methods

### Key resources table

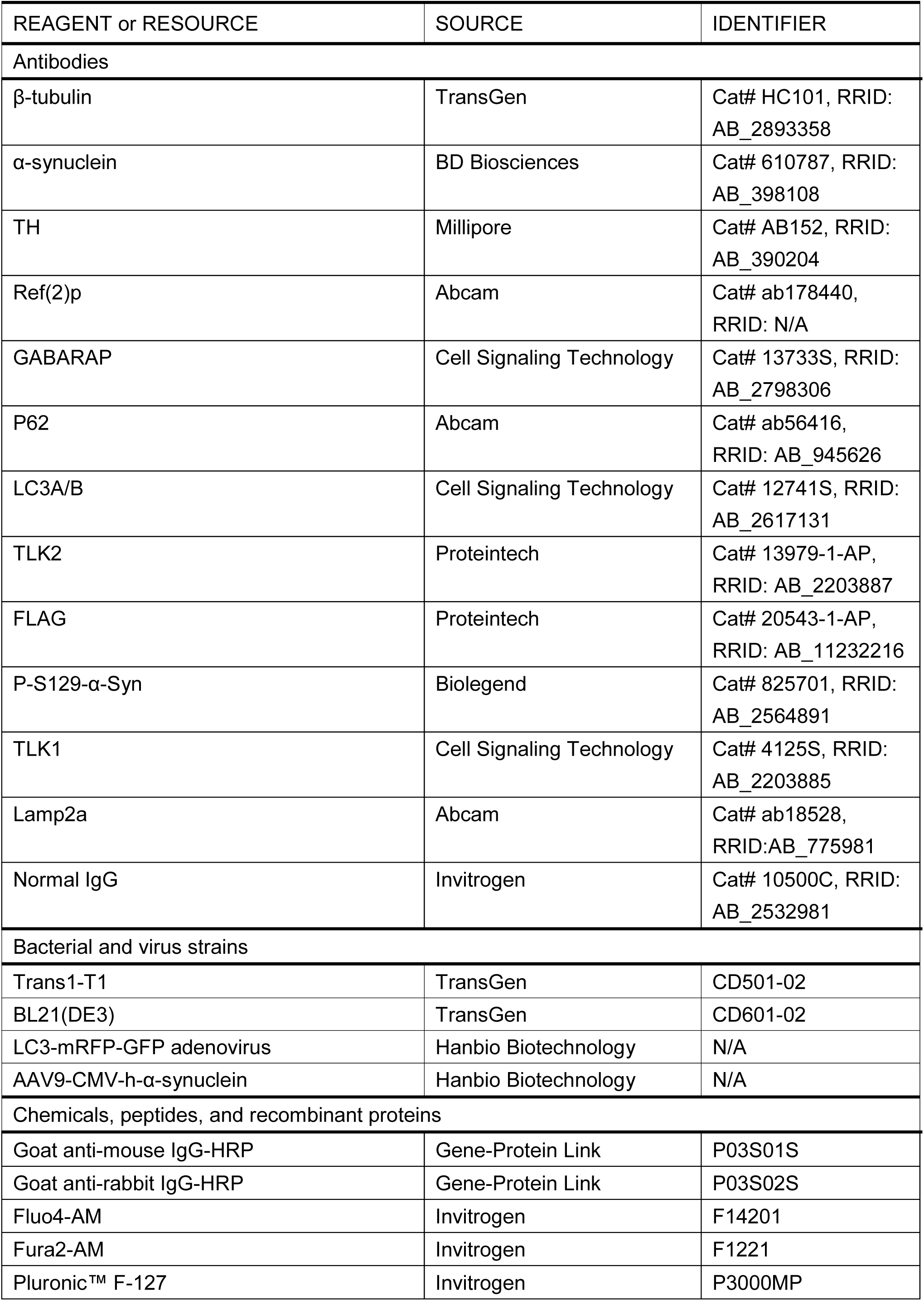
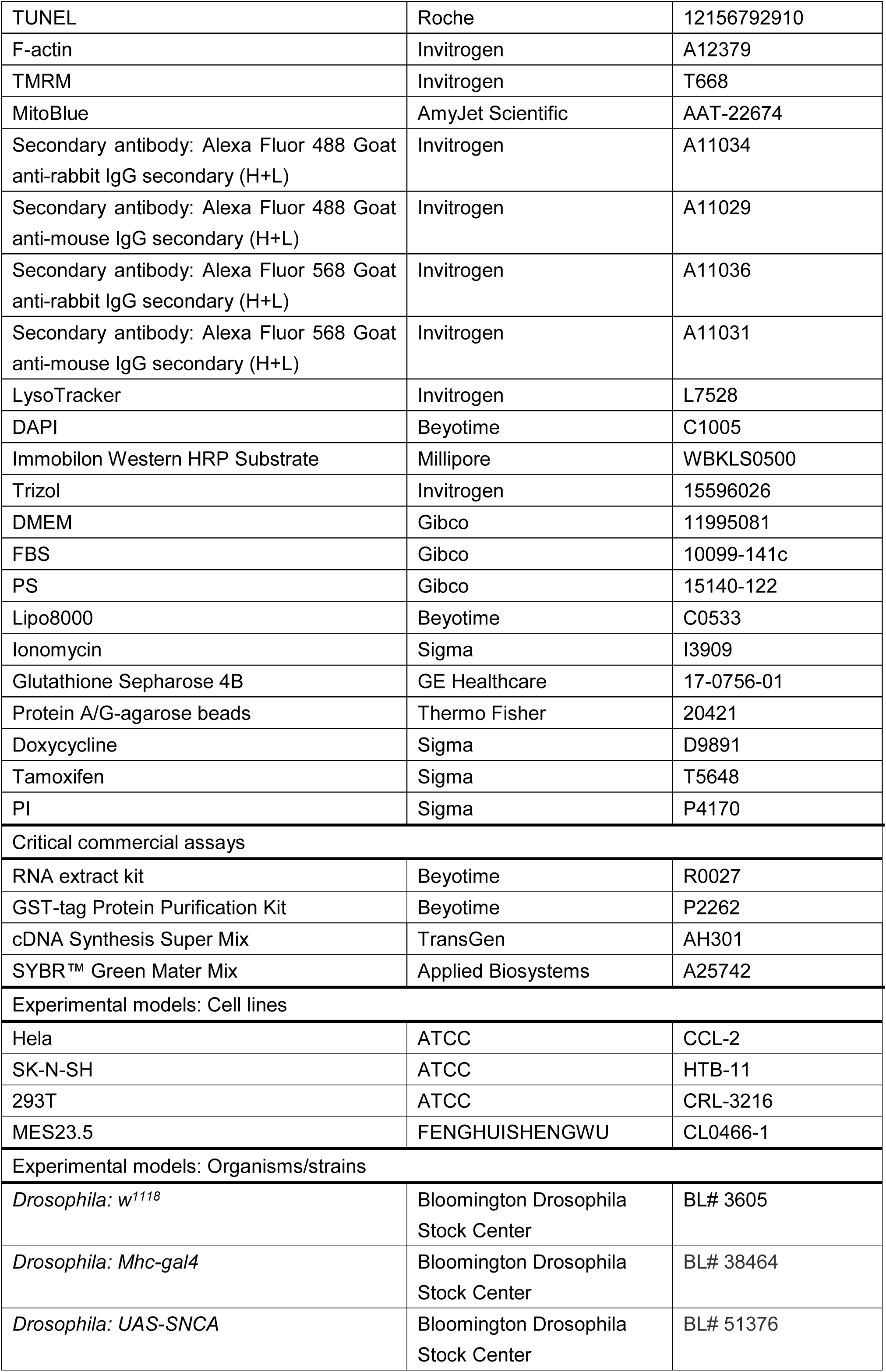
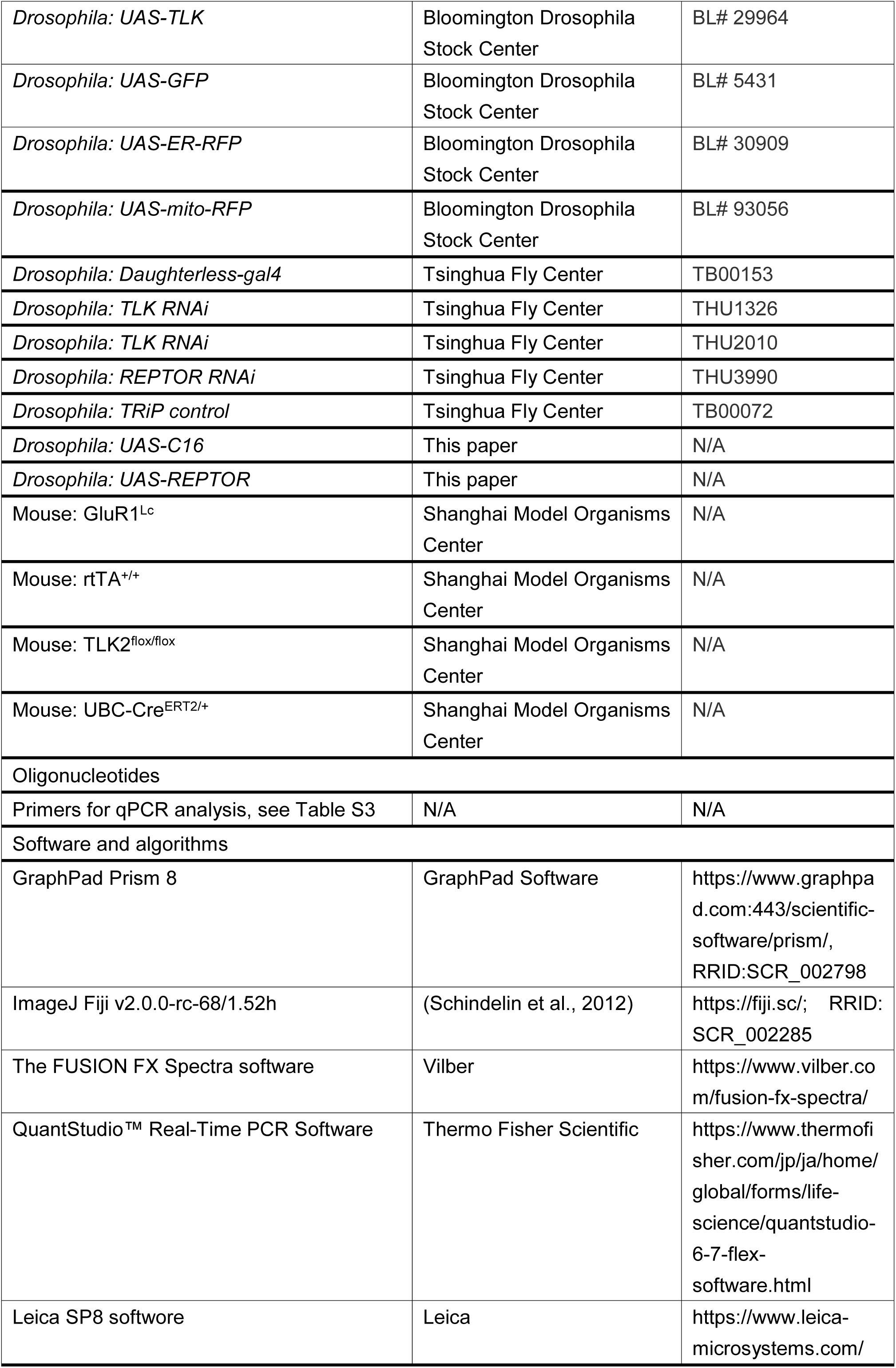

### *Drosophila* stocks and maintenance

*Drosophila* stocks were maintained under standard conditions at 25 ℃ on agar, cornmeal and yeast food. The flies included *w^11^*^18^*, Mhc-gal4, UAS-SNCA, UAS-TLK, UAS-GFP, UAS-ER-RFP* and *UAS-mito-RFP* were purchased from Bloomington *Drosophila* Stock Center; *Daughterless-gal4, TLK RNAi, REPTOR RNAi, TRiP control* and all other TRiP lines were obtained from Tsinghua Fly Center. The *UAS-C16* and *UAS-REPTOR* transgene were generated under the *w^11^*^18^ background by P-element insertion in our laboratory. For *DaGS>SNCA* flies, 20 flies were collected and placed in one tube at 1-3 days, 500 μM RU486 (Mifepristone; MedChemExpress #HY-13683) was mixed in the standard food, the fly vials were changed to new vials every two days. For *MC16* fly survival, 1-3 days-old flies were collected and transferred to new vials every two days. The dead flies were recorded until all flies were dead.

### Western blot

Tissues or cells were lysed in RIPA buffer (50 mM Tris, 150 mM NaCl, 1% Triton X-100, 1% sodium deoxycholate and 0.1% SDS) supplemented with protease inhibitor cocktail (Roche #11697498001) and PMSF (Beyotime #ST507) for 30 minutes on ice. Then the lysis was sonicated for 1 minute with 40 w 3 sec on / 3 sec off on ice and centrifugation at 12,000 rpm 4 ℃ for 15 minutes. Supernatants were collected and boiled with SDS loading buffer at 95 °C for 5 minutes. 20-30 μg protein from cell lysate or tissue homogenate were loaded on 12% SDS-PAGE gels. Transferring the protein from the gel to the PVDF membrane (Millipore #ISEQ00010). Membranes were blocked with 5% milk for one hour at room temperature and incubated with appropriate dilutions of primary antibody overnight at 4 °C. Washing the membrane 3 times for 5 minutes each with TBST (TBS containing 0.1 % Tween20), and then incubating with HRP secondary antibody (1:10000) for one hour at room temperature. Acquiring image using HRP substrate (Millipore #WBKLS0500) for chemiluminescence with VILBER Fusion FX SPECTRA equipment.

### Phos-tag Western blot

Mice brain tissues were quickly homogenized in ice-cold RIPA buffer supplemented with protease inhibitor cocktail, PMSF and phosphatase inhibitor (Beyotime #P1082) for 30 minutes on ice. Then the lysis was centrifugation at 12,000 rpm 4 ℃ for 15 minutes, supernatants were collected, part of the supernatants were treated with alkaline phosphatase (Beyotime #D7027) and boiled with SDS loading buffer at 95 °C for 5 minutes. 40 μmol/L MnCl_2_ and 20 μmol/L phos-tag (Wako #304-93521) were added to the SDS-PAGE (8 %). After the electrophoresis, the phos-tag gels were incubating with transfer buffer containing 5 mM EDTA for 20 minutes twice to remove the divalent cations ^49^. Washing with transfer buffer for 10 minutes once. Transferring the protein from the gel to the PVDF membrane. The next blocking and antibody incubation were the same as the normal western blot procedures.

### Immunofluorescence staining

Mice were anesthetized with pentobarbital sodium (80 mg/kg) and perfused with 0.9 % saline followed by 4 % paraformaldehyde (PFA). The brains were removed and placed in 4 % PFA overnight, then transferred to 10 %, 20 % and 30 % sucrose until the brains dropped to the bottom. The brains were embedded in OCT (SAKURA #4583) and cut into 10 μm thick coronal slices. The *Drosophila* brain or indirect flight muscle were dissected in HL3 buffer (70 mM NaCl, 5 mM KCl, 0.2 mM CaCl2, 20 mM MgCl2, 10 mM NaHCO3, 5 mM trehalose, 115 mM sucrose, and 5 mM HEPES). Brain slices, *Drosophila* tissues or cells were fixed with 4 % PFA for 15 minutes at room temperature, and then incubated three times for 5 minutes each with PBST (PBS containing 0.2% Triton X-100) for permeabilization, followed by blocking with 5 % BSA in PBST for 30 minutes at room temperature. Then incubation with primary antibody in the blocking solution overnight at 4 ℃. Washing with PBS three times for 5 minutes each with gentle shaking. Incubating with secondary antibody (1:200) for 1 hour (light protected) and repeat washing, optional staining with DAPI solution (Beyotime #C1005) for 5 minutes and rinsing with PBS. Mounting coverslip with mounting medium (abcam #ab104135). Acquiring images with Leica SP8 confocal microscope.

### Quantitative real-time PCR

Total RNAs were extracted according to the manufacturer’s protocol (Beyotime #R0027). 5 μg of total RNA were reversed using the TransScript® II First-Strand cDNA Synthesis SuperMix kit (TransGen #AH301) based on the manufacturer’s instructions. The final volume of qRT-PCR reaction mixer was 10 μl containing 5 μl PowerUp SYBR Green Master Mix (Applied Biosystems #A25742), 1 μl diluted cDNA sample (1:1), 0.5 μl forward primer, 0.5 μl reverse primer and 3 μl Nuclease-free water. The quantification of target genes was calculated by ΔCt method with Applied Biosystems QuantStudio™ Real-Time PCR System. Actin was used as a reference.

### Fluo4-AM staining

Fly brains or muscles were dissected in HL3 buffer, and incubated with 5 μM Fluo4-AM (Invitrogen #F14201) and 0.02 % Pluronic F-127 (Invitrogen #P3000MP) at 25 °C for 30 minutes in the dark. Then washing with HL3 buffer three times for 10 minutes each. Measuring fluorescence using Leica SP8 confocal microscope for excitation at 494 nm and emission at 516 nm ^50^.

### Fura2-AM staining

Fly indirect flight muscles were dissected in HL3 buffer, and incubated with 1 μM Fura2-AM (Invitrogen #F1221) and 0.02 % Pluronic F-127 (Invitrogen #P3000MP) at 25 °C for 30 minutes in the dark. Then washing with HL3 buffer three times for 10 minutes each. Measuring fluorescence using IONOPTIXBIOION-1.3 confocal microscope for excitation at 340 nm and 380 nm and the fluorescence intensities detected at ∼510 nm by WinFluor software.

### Histology of *Drosophila* adult thorax or brain

Drosophila indirect flight muscle (IFM) were dissected in HL3 buffer, GFP, mito-RFP and ER-RFP were imaged by Leica SP8 confocal microscope respectively. Fly tissues were incubated with 1 μM lysotracker (Invitrogen #L7528) at 25 °C for 10 minutes in the dark. Then washing with HL3 buffer three times for 10 minutes each. For TMRM (Invitrogen #T668) staining, the experimental procedures were similar to the lysotracker staining. For MitoBlue (AmyJet Scientific #AAT-22674) staining, Fly tissues were incubated with 10 μM MitoBlue at 37 °C for 60 minutes in the dark. Then washing with HL3 buffer three times for 10 minutes each.

### TUNEL staining

Fly indirect flight muscle (IFM) were dissected and fixed with 4 % PFA for 15 minutes at room temperature, and then incubated three times for 5 minutes each with PBST (PBS containing 0.2 % Triton X-100), followed by adding 10 μg/ml proteinase K solution diluted with PBS for 10 minutes at 56 °C. Washing with PBST three times for 5 minutes each. The TUNEL (Roche #12156792910) reaction mixture were incubated for 2 hours at 37 °C, and protected from light. Washing with PBST three times for 5 minutes each. Then F-actin (Invitrogen #A12379) were dissolved into DMSO to make a 1000x stock solution, diluting this stock solution 1:1000 in PBS and staining for one hour at room temperature, Washing with PBS three times for 5 minutes each. Acquiring images with Leica SP8 confocal microscope.

### Transmission electron microscopy

The fly muscles or cells were fixed with 2.5 % glutaraldehyde in 0.1 M PB buffer for 4 hours at 4 °C. Washing with 0.1 M PB 3 times for 10 minutes each. Then sample were prepared for imaging according to the standard protocol and observed using a transmission electron microscopy (HT7700).

### RNA sequencing

Total RNAs were extracted using Trizol (Invitrogen #15596026) reagent based on the manufacturer’s instructions, the integrity and quantity of RNA sample using agarose gel electrophoresis and NanoDrop ND-1000 (Thermo Scientific, USA). RNA sample were sequenced by Illumina HiSeq 2500 instrument, differentially expressed genes were analyzed.

### Generation of TLK1 and TLK2 knockout in Hela cells

Guide RNAs were designed by CRISPOR (http://crispor.tefor.net/crispor.py), TLK1 gRNA (5’-AAAGTATTGGGGGA-3’) was recognized the Exon5 and TLK2 gRNA (5’-ATATCTCTAGGCAACAGGAA-3’) was recognized the Exon12. gRNAs were cloned into Cas9 vector-PX459 and then transfected to the Hela cells with Lipo8000 (Beyotime #C0533). After 48 hours transfection, 3 μg/ml puromycin were added to the fresh medium, cells were cultured for 24 hours to eliminate the cells without recombinant plasmid. Then transfer the survival cells into a 96-well plate to expand single clones for approximately 3 weeks, genomic DNA were extracted and sequencing after PCR amplification using the following primers TLK1: F: TCACTTTGTCAGCGTGGTCA, R: CCACGGTCTGACTGCAAAAA; TLK2: F: CGTGTAGAGTAGTAATGGCTCCC, R: ACCAGGAGTGGCCAAAAGTC. TLK1 and TLK2 knock out cells were confirmed by western blot.

### LC3-mRFP-GFP adenovirus transfection

Hela cells were cultured in DMEM medium (Gibco #11995081) supplemented with 10 % FBS (Gibco #10099-141c), 1 % streptomycin and penicillin (Gibco #15140-122). When cell density was between 50 % and 70 % in confocal dish, LC3-mRFP-GFP adenovirus (Hanbio Biotechnology) were added to 1 ml medium at multiplicity of infection (MOI) of 10 ^51^. After 6-8 hours transfection, changing the medium and further culturing for 24 hours, cells were imaged with Leica SP8 confocal microscope.

### GST pull down assay

For TLK2 purification, TLK2 cDNA was cloned into the PGEX-6P-1 vector containing a GST tag. This construct was expressed into the BL21 chemically competent cells (TransGen # CD701), and the induced condition was 0.5 mM IPTG, 180 rpm rotation for 20 hours at 20 ℃. GST-TLK2 was purified by GST-tag Protein Purification Kit (Beyotime #P2262) according to the manufacturer’s protocol.

Hela cells were treated with 1 μM ionomycin (Sigma # I3909) for 3 hours when the cells density reached to 70 % confluence. Then cells were harvested and lysed in lysis buffer (10 mM Tris, 150mM NaCl, 2 mM EDTA, 0.5% TritonX-100). After centrifugation at 12,000 rpm 4 ℃ for 15 minutes, supernatants were collected. Mixing 3 μg GST-TLK2 purification protein and 200 μg cell lysate, rotation at 4 ℃ overnight. After rotation, GST-TLK2 were pulled down by Glutathione Sepharose 4B (GE Healthcare #17-0756-01) for 1 hour rotation at room temperature. Washing with lysis buffer 6 times at 2000 rpm 4 ℃ for 5 minutes each. Interactions between GST-TLK2 and beads were disrupted by 4x loading buffer at 95 ℃ for 5 minutes. The final results were obtained by SDS-PAGE, phos-tag SDS-PAGE, Coomassie Blue staining and mass spectrometry, or TLK2 protein were cut by PreScission Protease (Beyotime #P2302) overnight at 4 ℃.

### TLK2 kinase assay

Standard kinase assay was performed for 15 minutes at room temperature in 50 μl of kinase buffer (10 mM Tris pH 7.5, 50 mM KCl, 10 mM MgCl2, 1 mM DTT) supplemented with 1 μl ATP (0.2 mM) and with 1 μg TLK2 purification proteins and 20 μg substrates. MBP (Millipore #13–110), PGK1 or CREBRF were the substrates used for the kinase assays ^29^.

For CREBRF purification, CREBRF cDNA was cloned into the PGEX-4T-1 vector containing a GST tag. The induced condition was 0.5 mM IPTG, 200 rpm rotation overnight at 30 ℃. CREBRF protein was purified by GST-tag Protein Purification Kit (Beyotime #P2262) according to the manufacturer’s protocol.

### Co-immunoprecipitation

CREBRF-FLAG was transfected in the HEK293T cells for 48h. The cells were lysed in the lysis buffer (20 mM Tris pH7.5, 150 mM NaCl, 1% Triton X-100) supplemented with protease inhibitor cocktail and PMSF for 30 minutes on ice. After centrifugation at 12,000 rpm 4 ℃ for 15 minutes, supernatants were collected and incubated with 3 μg FLAG antibody (Proteintech #20543-1-AP) or control IgG (Invitrogen #10500C) rotating at 4 °C overnight. The protein A/G-agarose beads (Thermo Fisher #20421) were added to precipitate the complexes at 4 °C overnight. After washing with lysis buffer six times at 2000 rpm 4 ℃ for 5 minutes each, the agarose beads were boiled in a 4x SDS loading buffer. The results were obtained by western blot as described above.

### GluR1^Lc^ transgenic mice

All the mice were housed in the animal center of Capital Medical University. The animal studies were approved by the Institutional Animal Care and Use Committee, Capital Medical University, Beijing, China (approved number: AEEI-2015-156). GluR1^Lc^ mice were purchased from Shanghai Model Organisms Center. The SA-polyA-Insulator-Insulator-TRE-miniCMVpromoter-GluR1(Mut)-wpre-pA-Insulator-Insulator-FRT-PGK-Neo-pA-FRT was inserted at the Gt(ROSA)26Sor (ENSMUSG00000086429) locus by In-Fusion cloning. The GluR1^Lc^ ^m/m^ (ENSG00000155511) transgenic mice were crossed with rtTA^+/+^ mice, doxycycline (Sigma #D9891) was given in drinking water to induce GluR1^Lc^ expression.

### TLK2 knockout mice

TLK2 knockout mice were also purchased from Shanghai Model Organisms Center. The Cre-lox system was used for the deletion of TLK2 gene (ENSG00000146872). TLK2^flox/flox^ mice were generated by CRISPR-Cas9 technology which targeted exon4. After mating them with UBC-Cre^ERT2/+^ mice, TLK2 genes could be knocked out ubiquitously by tamoxifen (Sigma #T5648) via intraperitoneal injection in 75 mg/kg body weight once every 24 hours for a total of 5 consecutive days. After the conditional knockout of TLK2, the mice appeared normal, including appearance, weight gain, mobility, and eating and drinking.

### Stereotactic injection of virus

Male, 8-10 weeks old, C57BL/6 mice were anesthetized with pentobarbital sodium (80 mg/kg) and placed in a stereotaxic frame (RWD, china). AAV9-CMV-h-α-synuclein (Hanbio Biotechnology) was injected into in the appropriate location (AP:-3.0 mm; ML:-1.3 mm; DV:-4.5 mm) for the substantia nigra (SN) at the right of the skull ^52^. 1 μl of AAV9-CMV-h-α-synuclein (concentration: 1.5 x 10^12^ vg/ml) or PBS were injected at a rate of 0.1 μl/min with a 10 μl Hamilton syringe. The needle was left in the place for a further 5 minutes after injection to avoid leakage. After the surgery, mice were placed on a warm electric blanket for recovery.

### Mice behavioral tests

In the open field test, mice were acclimated to the experimental room in their home cages for 30 minutes to minimize stress and then were placed into the open field cage (50 × 50 × 50 cm). After 1 minute of adaptation, the mice were allowed to explore the arena for 5 minutes. After the 5 minutes recording by SMART 3.0 software (RWD, china), mice were returned to their home cages and the arena was cleaned with 70 % ethanol ^53^.

In the rotarod test, mice were placed on an accelerating rotarod at speeds between 5 and 45 rpm for a maximum of 5 minutes. The animals were tested for 3 days with 2 trials per day, the latency to fall was recorded and the last 2 trials of the 3^rd^ day were analyzed for locomotor abilities ^54^.

### Statistical analyses

For mouse and rat experiments, the animal labeling was mutual blinded among the experimenters. Due to unknown standard deviation, we did not determine sample size before the experiments, without exclude any animal. Results were presented as mean ± SD. Statistical significance for comparisons between data sets was performed with an unpaired t-test between two groups. For comparison of more than two groups, significance was determined using a one-way ANOVA with Tukey’s post hoc test. Kaplan-Meier tests was used under the log-rank algorithm for survival analysis. Statistical analyses were performed using GraphPad Prism software. P value of < 0.05 was considered significant (* P < 0.05, ** P < 0.01, *** P < 0.001 and ****p < 0.0001).

---

**Figure S1.**
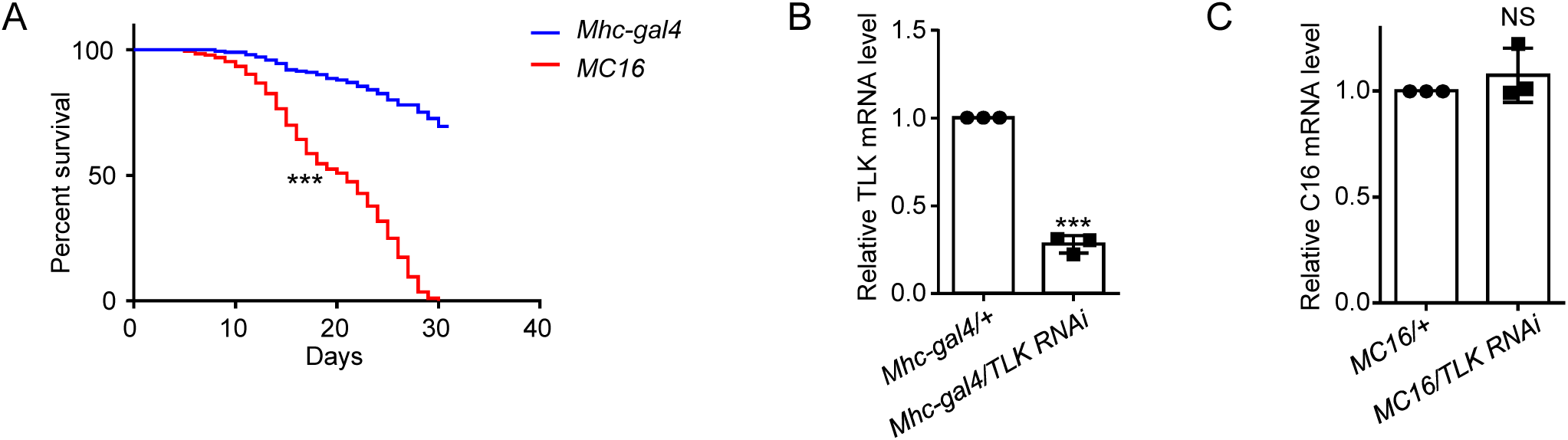
Characterization of *MC16* flies. (A) The lifespan changes of *Mhc-Gal4* (wild type control) and *MC16* flies. 80-100 flies were tested for each trial (N=3). (B) Quantitative RT-PCR to analyze TLK mRNA expression from *Mhc-Gal4/+* and *Mhc-Gal4/TLK RNAi* flies. 20 flies were tested for each trial (N=3). (C) Quantitative RT-PCR to analyze C16 mRNA level from *MC16/+* and *MC16/TLK RNAi* flies. 20 flies were tested for each trial (N=3).

**Figure S2.**
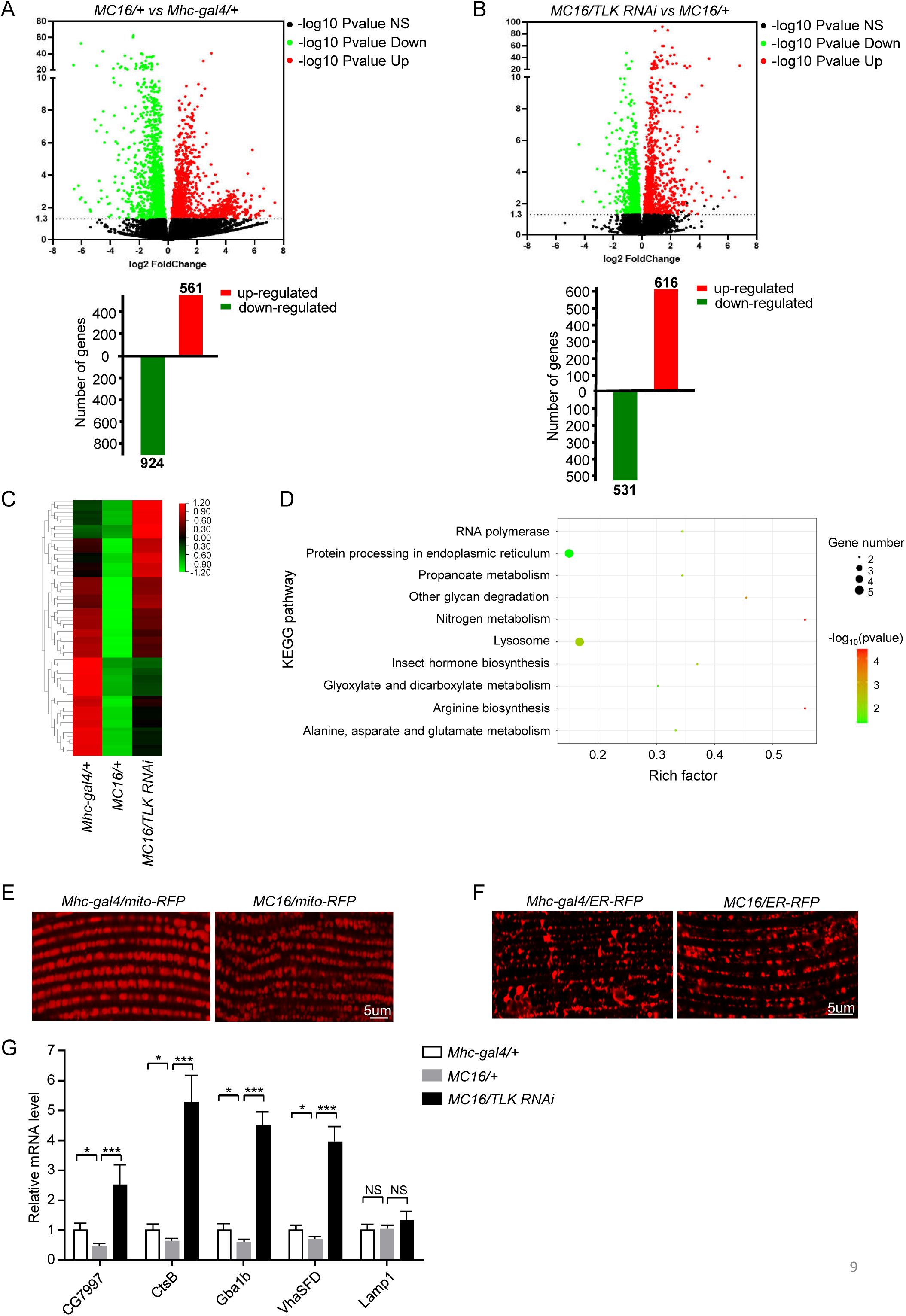
Analysis of global transcriptional changes in *Mhc-Gal4/+*, *MC16/+* and *MC16/TLK RNAi* flies; and mitochondria and ER morphological alternations in *MC16* flies. (A and B) The volcano plot of gene expression difference between groups of flies as indicated on the graph with p-value < 0.05. The red dots indicate upregulated genes, green dots indicate the downregulated genes, 561 genes upregulated and 924 genes downregulated in the *MC16* flies. Compared the *MC16* flies, TLK RNAi under *MC16* background results in 616 genes upregulated and 531 genes downregulated. Three independent samples were collected for each phenotype. The full RNAseq data are deposited in the supplementary source data sets. (C and D) The heat map of RNA sequencing result. The overlapping genes of the downregulated genes in *MC16* and the upregulated genes in *MC16/TLK RNAi* were analyzed by KEGG pathways. The enriched pathways are indicated on the graph. (E) Live imaging of the mitochondrial morphology of the indirect flight muscle. The mitochondria are labeled with *UAS-mitoRFP* driven by *Mhc-Gal4*. Scale bar, 5 µm. N=10. (F) Live imaging of the ER morphology of the indirect flight muscle. The ER is labeled with *UAS-ER-RFP* driven by *Mhc-Gal4*. Scale bar, 5 µm. N=10. (G) Quantitative RT-PCR analysis of five lysosomal genes mRNA expression from *Mhc-gal4/+*, *MC16/+* and *MC16/TLK RNAi* flies. 20 flies were tested for each trial (N=3).

**Figure S3.**
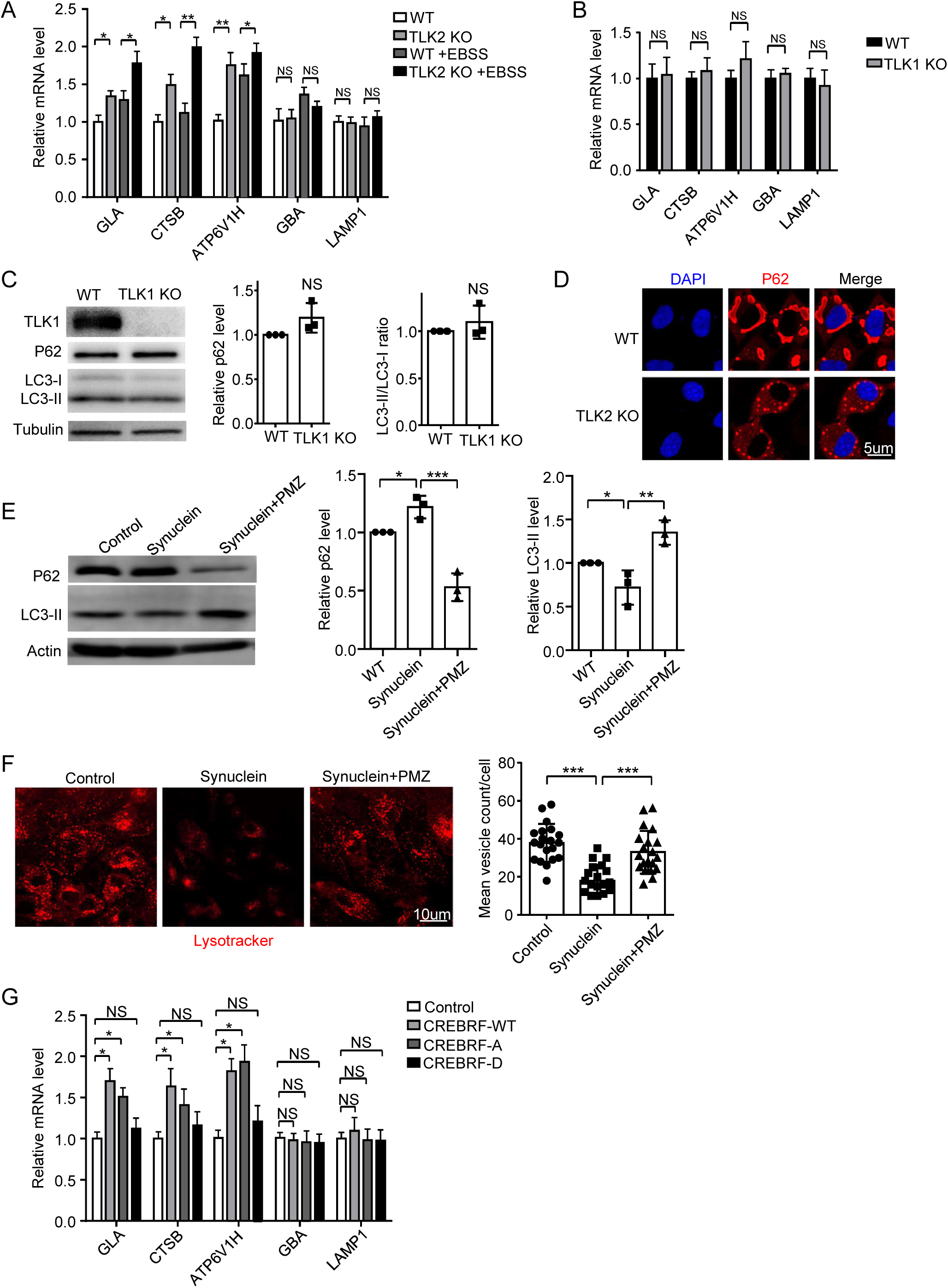
TLK1 and TLK2 KO effect on transcriptional level of lysosomal genes and autophagy-lysosome function. (A) Quantitative RT-PCR analysis of five lysosomal genes mRNA expression from Hela WT, TLK2 KO, WT+ starved in Earle’s starvation buffer (EBSS) and TLK2 KO+EBSS cells. EBSS is a powerful inducer of autophagy in cultured cells. N=3. (B) Quantitative RT-PCR analysis of five lysosomal genes mRNA level from Hela WT and TLK1 KO cells. N=3. (C) Western blot of the extracts from Hela WT and TLK1 KO cells. N=3. Quantification of P62 level and LC3-II/LC3-I ratio. (D) Immunostaining images of the P62 protein (red) in the P62 overexpression Hela cells, nuclei were stained with DAPI (blue). Scale bar, 5 µm. N=3. (E) Western blot of the extracts from the α-synuclein overexpression, or α-synuclein overexpression plus PMZ (TLK kinase inhibitor) treatment, or control cells. N=3. α-synuclein, P62 and LC3-II/LC3-I ratio is quantified. (F) Lysotracker (red) staining in cells overexpressed α-synuclein, or α-synuclein overexpression plus PMZ treatment, or the control cells. Scale bar, 10 µm. N=20. Quantification of the number of lysotracker positive vesicles. (G) Quantitative RT-PCR analysis of five lysosomal genes mRNA level from CREBRF-WT (normal), CREBRF-A (dephosphorylation), CREBRF-D (phosphorylation) and control cells. N=3.

**Figure S4.**
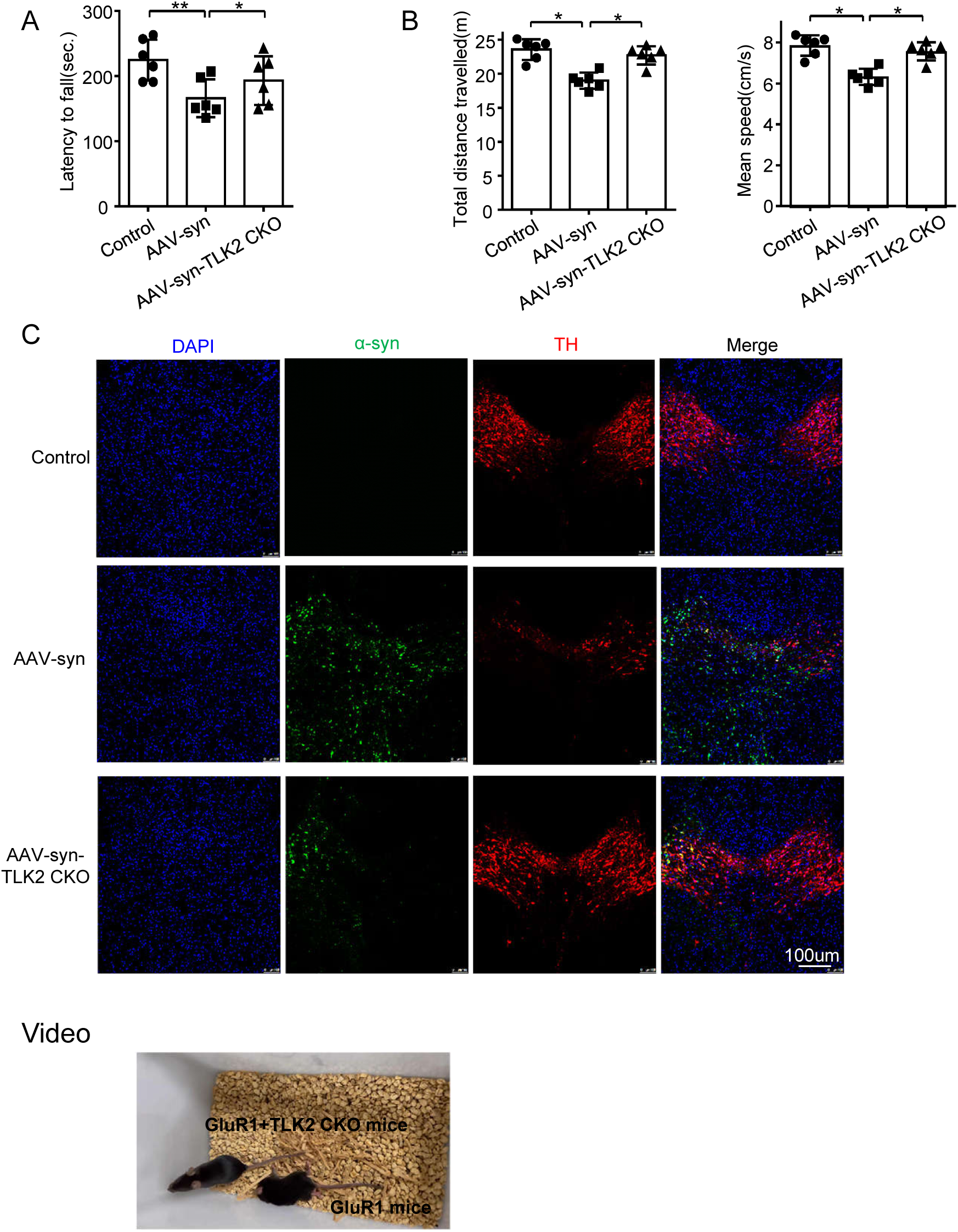
Effect of TLK2 KO on α-synuclein induced DA neuron loss and lethality of GluR1^Lc^ mice. (A) Rotarod behavioral tests of AAV-syn, AAV-syn+TLK2 CKO, and control mice. N=6 for each genotype. Quantification of the time of the latency to fall; (B) Open field behavioral tests of AAV-syn, AAV-syn+TLK2 CKO, and control mice. N=6 for each genotype. Quantification of the total distance and mean speed for 5 minutes in the field; (C) Immunostaining images of the TH neurons (red) in the SNc, nuclei were stained with DAPI (blue), α-synuclein were accompany with GFP fluorescent protein (green). Scale bar, 100 µm. N=4. (Supplementary video) The video displayed the mobility of GluR1^Lc^ and GluR1^Lc^ / TLK2 CKO mice after 5 days of dox treatment. GluR1^Lc^ mice showed defective mobility; whereas GluR1^Lc^ / TLK2 CKO mice displayed better mobility. N=6.

## Notes

### Competing Interest Statement

The authors have declared no competing interest.

